# Neural Network Organization for Courtship Song Feature Detection in *Drosophila*

**DOI:** 10.1101/2020.10.08.332148

**Authors:** Christa A. Baker, Claire McKellar, Aljoscha Nern, Sven Dorkenwald, Diego A. Pacheco, Rich Pang, Nils Eckstein, Jan Funke, Barry J. Dickson, Mala Murthy

## Abstract

Animals communicate using sounds in a wide range of contexts, and auditory systems must encode behaviorally relevant acoustic features to drive appropriate reactions. How feature detection emerges along auditory pathways has been difficult to solve due to challenges in mapping the underlying circuits and characterizing responses to behaviorally relevant features. Here, we study auditory activity in the *Drosophila melanogaster* brain and investigate feature selectivity for the two main modes of fly courtship song, sinusoids and pulse trains. We identify 24 new cell types of the intermediate layers of the auditory pathway, and using a new connectomic resource, FlyWire, we map all synaptic connections between these cell types, in addition to connections to known early and higher-order auditory neurons - this represents the first map of the auditory pathway. We additionally determine the sign (excitatory or inhibitory) of most synapses in this auditory connectome. We find that auditory neurons display a continuum of preferences for courtship song modes, and that neurons with different song mode preferences are highly interconnected in a network that lacks hierarchical structure. Among this network, frequency tuning is centered on the range of frequencies present in song, whereas pulse rate tuning extends to rates outside of song, suggesting that these neurons form a basis set for downstream processing. Our study provides new insights into the organization of auditory coding within the *Drosophila* brain.

## INTRODUCTION

Sounds are an integral part of the social life of animals, and are also critical for mate choice, finding food, caring for young, and avoiding harm. Accordingly, the brains of animals have evolved to recognize behaviorally salient acoustic signals. For instance, courtship songs often contain information about sender status and species, and receivers must decode this information by analyzing patterns within songs (Akre et al., 2011; Baker et al., 2019; Hedwig, 2016; Nieder and Mooney, 2020). Several species that use sound to communicate produce songs comprising multiple acoustic types, often referred to as syllables or modes (Behr and von Helversen, 2004; Holy and Guo, 2005; Wohlgemuth et al., 2010). While prior work has examined where selectivity for conspecific sounds emerges across a variety of systems including songbirds (Moore and Woolley, 2019), primates (Romanski and Averbeck, 2009), and mice (Roberts and Portfors, 2015), ***how*** this selectivity arises remains unknown. While a hypothesized circuit for recognizing stereotyped conspecific songs has been described in crickets (Clemens et al., 2020; Schöneich et al., 2015), synaptic connectivity between neurons in the circuit has not yet been determined, and cricket songs comprise only a single mode. Ultimately, understanding how auditory systems establish selectivity for conspecific sounds requires linking information on responses to different song types with neuronal connectivity, a significant challenge in larger brains. Here we focus on *Drosophila melanogaster*, a species with a compact brain and both genetic and connectomic tools for circuit dissection.

*D. melanogaster* males alternate between pulse and sine song (Fig. 1A; (Arthur et al., 2013; Bennet-Clark and Ewing, 1967)), dependent on sensory feedback they receive from a female, to compose song bouts (Coen et al., 2014). Receptive females reduce locomotor speed in response to features within each mode, from the frequency of the individual elements to longer timescale patterns such as the duration of each mode (Clemens et al., 2015; Deutsch et al., 2019). In general, both males and females respond selectively to features and timescales present in conspecific song, but how tuning for these features arises along the auditory pathway is not yet known.

**Figure 1.**
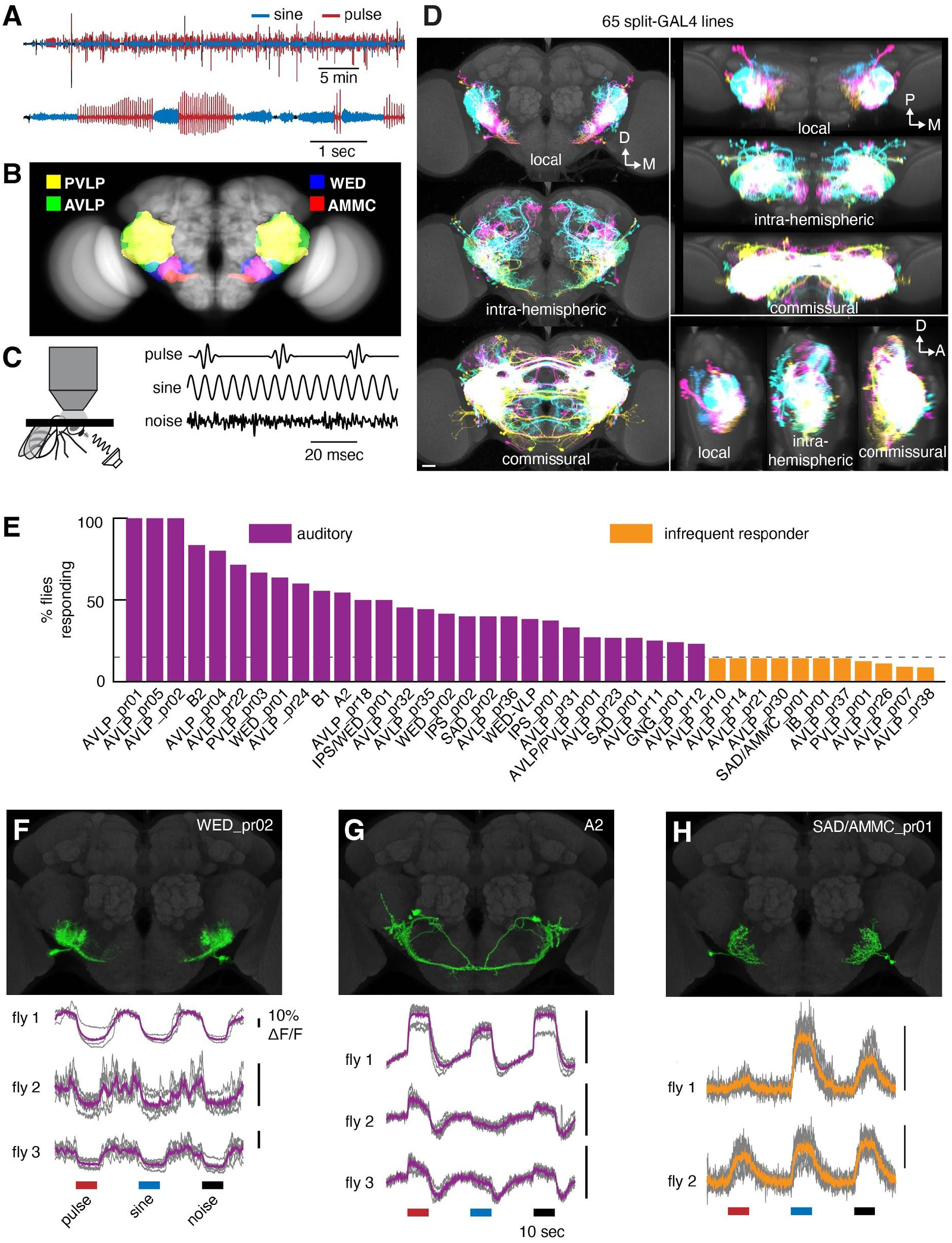
Anatomic and functional screen for auditory neurons. A) Microphone recording from a single wild-type (CS-Tully strain) male fly paired with a virgin female. The top trace shows song over 30 minutes, and the bottom trace shows a close-up of song bouts consisting of switches between the pulse and sine song modes. B) Primary auditory neurons called Johnston Organ neurons in the antenna project to the antennal mechanosensory and motor center (AMMC) in the central brain. Auditory information is then routed to downstream areas including the wedge (WED), anterior ventrolateral protocerebrum (AVLP), and posterior ventrolateral protocerebrum (PVLP). C) Schematic showing two-photon calcium imaging set-up with sound delivered to the aristae (left) and calibrated, synthetic acoustic stimuli used to search for auditory responses (right). One second of each stimulus is shown. D) Overlaid images of the split-GAL4 collection’s local interneurons, intra-hemispheric projection neurons, or commissural neurons, segmented from aligned images of the split GAL4 collection and shown as maximum projections from the front (left), top (top right), and side (bottom right). D: dorsal; M: medial; P: posterior; A: anterior. Scale bar: 25 microns. E) Percentage of imaged flies from each cell class with auditory responses. We defined auditory cell classes as those in which >15% (dotted line) of imaged flies responded to the pulse, sine, or noise stimuli. If at least 1 fly but fewer than 15% of imaged flies responded, we termed the cell class an ‘infrequent responder’. If no flies responded (out of 4-6 total flies), we termed the cell class ‘non-auditory’. Numbers of flies imaged ranged from 4-17 for auditory cell classes, and from 7-23 for infrequently responding cell classes. F-H) Calcium responses to pulse, sine, and noise stimuli from three cell classes. In the calcium traces, each trial is shown in grey and the mean across trials is shown in purple for the auditory cell classes in F-G and in orange for the infrequently responding cell class in H.

Sound detection begins at the *Drosophila* antenna. Johnston’s organ neurons (JONs) within the antennal second segment detect sound-evoked vibrations of the arista (Albert et al., 2007). Similar to vertebrate auditory systems, the fly auditory system parses sound frequency at the periphery. JONs appear to be coarsely tuned to frequency and project roughly tonotopically to the antennal mechanosensory motor center (AMMC) in the brain (Patella and Wilson, 2018) - tonotopy appears to be strengthened in downstream areas, like the wedge (WED) (Patella and Wilson, 2018) and ventrolateral protocerebrum (VLP) (Pacheco et al., 2020) (Fig. 1B). The auditory responses of a handful of AMMC neurons have been characterized (Azevedo and Wilson, 2017; Kamikouchi et al., 2009; Lai et al., 2012; Tootoonian et al., 2012), and one of these, called B1, is thought to be part of a major pathway for song processing, as silencing these neurons impacts song-evoked behaviors in both sexes (Vaughan et al., 2014; Yamada et al., 2018). Only a handful of additional auditory neurons have been identified (Clemens et al., 2015; Deutsch et al., 2019; Lai et al., 2012; Wang et al., 2020b; Zhou et al., 2015), and previous studies have not systematically examined, at the level of individual cell types, preference for the two main modes of song, pulse and sine. We therefore do not know how early auditory circuits are organized - whether sine and pulse processing are separate or intermingled, and whether tuning for song features is sharpened along the pathway in a hierarchical fashion.

In downstream brain areas, beyond the aforementioned mechanosensory regions, we now know of at least four cell types with pulse song feature preference. pC2l neurons in both males and females respond selectively to several distinct features that define pulse song, from the frequency of individual pulses to the length of pulse trains, but they respond only weakly to sine song (Deutsch et al., 2019). vpoDN/pMN2 neurons, and their inputs vpoEN and vpoIN, also respond more strongly to pulse vs. sine song (Wang et al., 2020b). Neurons selective for sine song features have not yet been identified, although pan-neuronal imaging suggests responses to sine song dominate throughout the brain (Pacheco et al. 2020). Understanding how preference for pulse vs. sine song emerges along the *Drosophila* auditory pathway is critical for revealing how the fly brain encodes and decodes courtship song, since both modes are known to affect female behavior during courtship (Clemens et al., 2018a; Deutsch et al., 2019; Schilcher, 1976).

We address these outstanding issues here, by uncovering the network organization underlying song processing in the *Drosophila* brain. We recorded auditory responses from a collection of split-GAL4 lines, and found 24 novel auditory cell types, as well as new lines for known auditory neurons. We systematically characterize each cell’s preference for sine vs. pulse song, in addition to tuning for sine frequency and pulse rate, discovering a continuum of song mode preferences among auditory neurons. Using FlyWire, a whole brain connectomic resource (Dorkenwald et al., 2020), we mapped synaptic connectivity (in addition to the sign of connections, whether inhibitory or excitatory) among auditory neurons. Rather than being organized into separate streams for sine and pulse processing, we discovered a complex neural network architecture with dense interconnectivity between cell types with different song mode tunings. In addition, we found that preference for sine and pulse song arises early in the pathway, suggesting that interconnectivity is used to generate the continuum of song mode preferences.

## RESULTS

### Identifying cell types of *Drosophila* auditory circuits

To identify new cell types downstream of known early auditory neurons (Dorkenwald et al., 2020), we focused on three of the five brain areas to which second-order mechanosensory neurons in the AMMC send the majority of their projections (Matsuo et al., 2016): the wedge (WED), anterior ventrolateral protocerebrum (AVLP), and posterior ventrolateral protocerebrum (PVLP) (Fig. 1B). Our general approach was to create sparse and specific split-GAL4 lines (Luan et al., 2006; Tuthill et al., 2013) targeting neurons (hereafter referred to as WED/VLP neurons) in any of these three areas and then to screen the most promising lines for auditory responses; our goal was not to create a line for every cell type in the WED/VLP (of which over a thousand are estimated to exist (Scheffer et al., 2020)), but to identify a wide variety of new auditory neuron types. In total, we examined the expression of 1041 split-GAL4 lines, generated stable lines for 117 with the sparsest expression in the brain, and selected 65 for functional recordings. These 65 lines target at least 50 WED/VLP cell types that include local, intra-hemispheric projection, and commissural neurons (Fig. 1D, Supp. Fig. 1).

To reveal cell type diversity within each split-GAL4 line, we collected multicolor flip-out (MCFO) images (see Methods) (Supp. Fig. 2). Most split-GAL4 lines contained morphologically similar neurons, with some heterogeneity within particular lines. As recent work on the *Drosophila* hemibrain has shown that even morphologically similar neuron types may have different presynaptic and postsynaptic partners (Scheffer et al., 2020), we were unable to disambiguate further the individual neuronal subtypes within each line on morphological criteria alone, and consider the WED/VLP neurons labeled in each line a morphologically similar ‘cell class’. We named each cell class by identifying the neuropil with the highest density of neural processes (Supp. Fig. 3; Table 2). Hereafter, we refer to the neurons imaged in each line by these names, except for neurons previously named and identified as auditory (i.e., A2, B1, B2, and WED-VLP neurons) (Table 2) (Dorkenwald et al., 2020; Kamikouchi et al., 2009; Lai et al., 2012). To clarify which group of cells we targeted in each line for calcium imaging, we digitally segmented WED/VLP neurons from the broader expression pattern of each line (Fig. 1F-H and Supp. Fig. 4).

We tested each WED/VLP cell type for auditory responses using GCaMP6s (Chen et al., 2013) and three stimulus categories: pulse, sine and broadband noise (Fig. 1C; see Methods for stimulus details). Pulse and sine constitute the two major modes of *D. melanogaster* courtship songs, and the noise stimulus was included to identify responses to acoustic features outside of the range of parameters in song. Of the 65 lines screened, we found consistent auditory responses in 28 (43%) (Fig. 1E), including four cell classes representing previously known auditory neurons: A2, B1, B2, and WED-VLP (Fig. 2; Table 2). An additional 11 cell classes (17%) were classified as infrequent responders because responses occurred in fewer than 15% of imaged flies (Fig. 1E). The remaining 26 cell classes did not show responses to acoustic stimuli.

**Figure 2.**
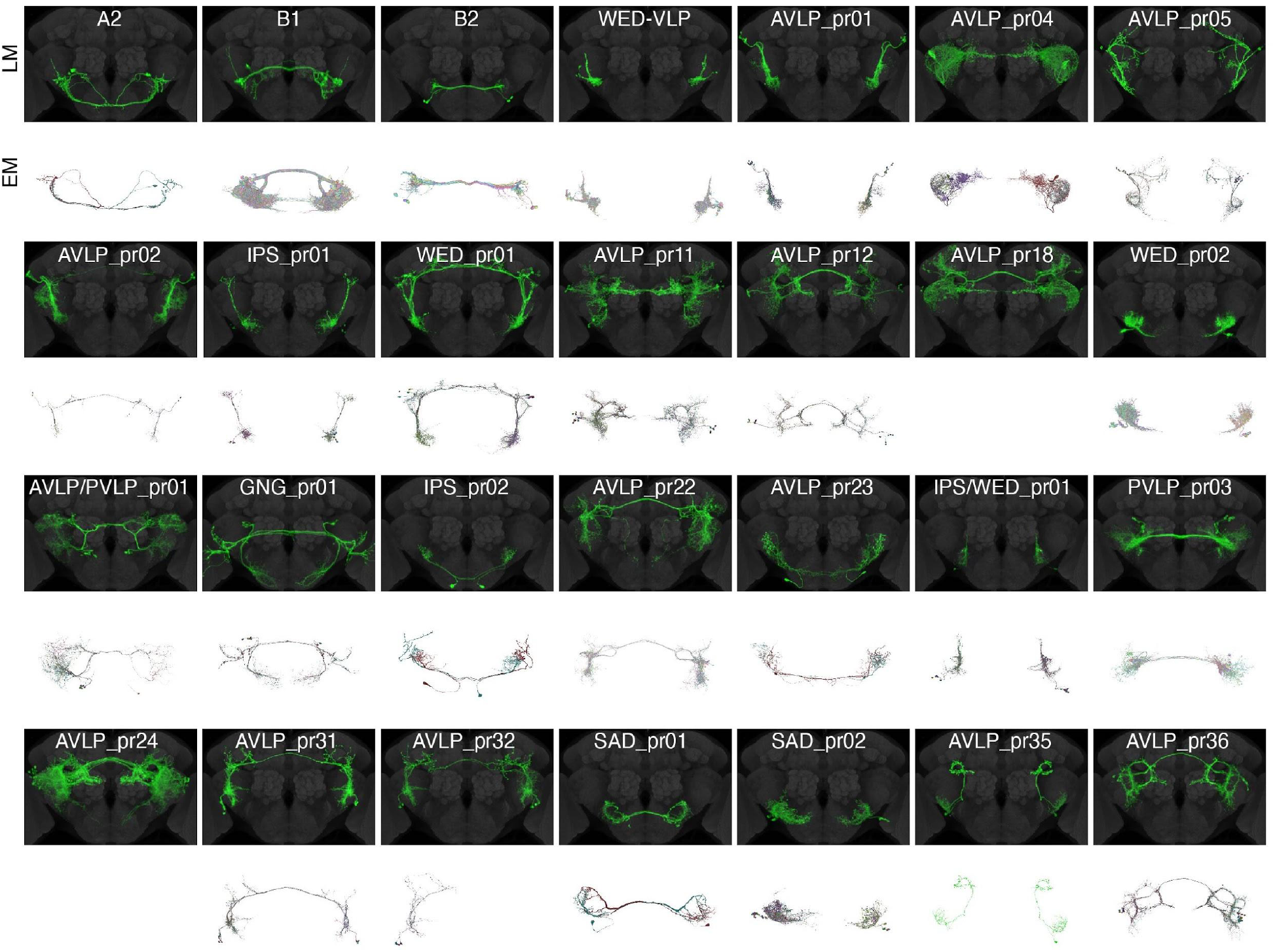
Light microscopic (LM) and electron microscopic (EM) images of auditory WED/VLP cell types. Aligned central brains with expression patterns of WED/VLP neuron classes digitally segmented (see Supp. Fig. 4). Only those cell classes with auditory responses are shown. Gray: nc82. Below each brain expression pattern are the EM reconstructions (identified and proofread in FlyWire.ai) corresponding to each cell class. There was insufficient information in the split-GAL4 and stochastic labeling expression patterns to resolve the EM reconstructions representing two cell classes (AVLP_pr18 and AVLP_pr24). EM reconstructions representing cell type AVLP_pr32 were only found in one hemisphere. AVLP_pr01 and AVLP_pr02 share morphological similarities with vpoINs, which provide sound-evoked inhibition onto descending neurons called vpoDNs that contribute to vaginal plate opening (Wang et al., 2020b). Based on both FlyWire and hemibrain connections, vpoINs (defined as inputs to vpoDN with morphology consistent with vpoINs) consist of two subtypes: one with a commissure, and one with a medial projection that does not cross the midline. This leads us to conclude that AVLP_pr02 are likely the commissural vpoINs, but AVLP_pr01 are a cell type independent from vpoINs.

We used the new FlyWire resource to find the auditory WED/VLP neurons in both hemispheres within an electron microscopic (EM) volume of an entire female brain (Fig. 2; see Methods) (Dorkenwald et al., 2020; Zheng et al., 2018). In most cases, we found clear morphological matches between light microscopic (LM) images and EM reconstructions, but in two cases (AVLP_pr18 and AVLP_pr24) we were not able to resolve the EM reconstructions belonging to each cell class based on the available LM images. We use these EM data below to examine connectivity between auditory cell types.

Some auditory neurons were previously classified as lateral horn neurons, but not known to be auditory (e.g., AVLP_pr05 resembles AVLP-PN1 and IPS_pr01 resembles WED-PN1) (Dolan et al., 2019), while others have morphologies similar to neurons that process other mechanosensory stimuli, such as wind (e.g., WED_pr01 is distinct from but morphologically similar to the WPN neurons that encode wind direction (Suver et al., 2019) and to the WPN_B_s, whose function has not yet been characterized (Coates et al., 2020)). In sum, our screen identified a total of 35 new neuron types with auditory activity. In the sections that follow, we investigate song feature tuning for these neurons, the diversity of their responses, and their synaptic connectivity.

### A continuum of preferences for courtship song modes among WED/VLP neurons

Auditory neurons tested for song spectrotemporal selectivity so far have been found to be pulse-preferring (Deutsch et al., 2019; Wang et al., 2020b; Zhou et al., 2015), but pan-neuronal imaging suggests there are a greater number of sine-preferring neurons throughout the central brain (Pacheco et al., 2020). We aimed to identify such elusive cell types in our screen. We found that the auditory responses of WED/VLP neurons consisted of increases/decreases in fluorescence indicative of net excitation, net inhibition, or a combination (Fig. 1F-H, 3A-C). To understand how these responses contribute to the encoding of courtship songs, we used the integral of the fluorescence trace during the stimulus to determine preference for pulse vs. sine stimuli in each fly imaged (Fig. 3A, Supp. Fig. 5A-B, and Methods). The effectiveness of this approach is supported by results from pan-neuronal imaging of auditory responses (Pacheco et al., 2020): when stimuli were morphed from a sine wave to a pulse train by varying the amplitude of a low-frequency envelope convolved with the stimulus, regions of interest in the WED and VLP regions preferred either the sinusoidal or the pulsatile stimulus, but not intermediate stimuli.

**Figure 3.**
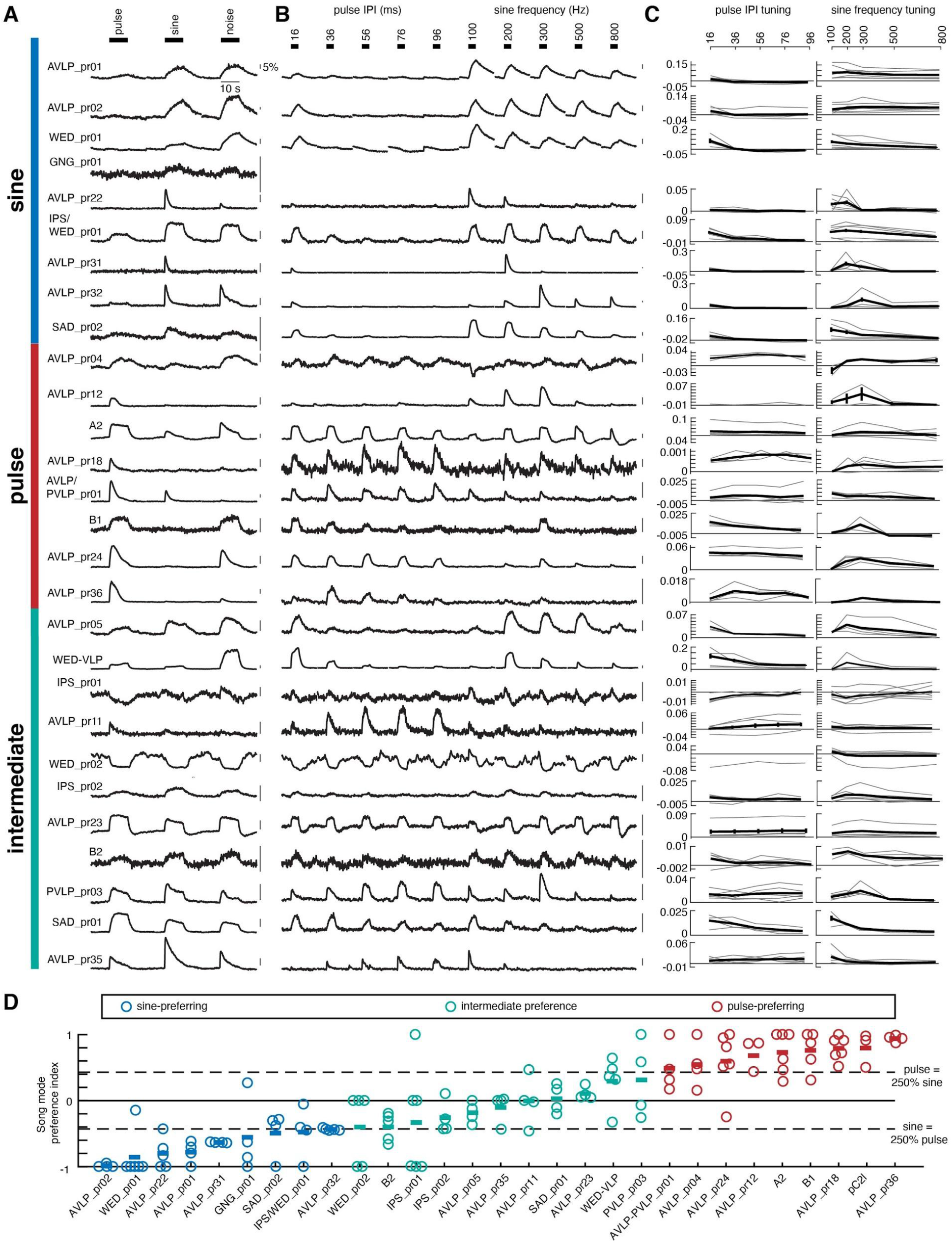
Auditory WED/VLP neurons show a continuum of preferences for sine and pulse song modes. A) Trial-averaged representative calcium traces for a single fly from each cell class in response to pulse, sine, and noise stimuli. Vertical colored bars indicate the pulse-vs. sine-preference of each cell class given in (D). B) Trial-averaged representative calcium traces in response to pulse rate (interpulse interval (IPI)) and sine frequency stimuli. Responses in (A) and (B) may come from the same or different flies. C) Pulse rate (left) and sine frequency (right) tuning curves. The tuning curves from individual flies are shown in grey, and the average across flies is shown in black. Error bars report standard error. D) The integrals of responses to pulse and sine were used to calculate a song mode preference index, which ranges from -1 (strongest sine preference) to 1 (strongest pulse preference) (see Methods; Supp. Fig. 5A-B). To identify sine-preferring cell types, we required the mean preference index across flies to be below -0.43, which corresponds to a sine response that is at least 250% that of pulse. To identify pulse-preferring cell types, we required the mean preference index across flies to be above -0.43, which corresponds to a pulse response that is at least 250% that of sine. All other cell types were classified as having intermediate song mode preference. The song mode preference index for pC2l was calculated using data from a previous study (Deutsch et al., 2019).

We quantified each cell type’s song mode preference by calculating a preference index (see Methods) that ranges from -1 (strongest sine preference) to +1 (strongest pulse preference). For a given WED/VLP neuron to be classified as pulse-or sine-preferring, we reasoned that the preferred song mode response should be at least 150% greater than the response to the other mode, which corresponds to a mean preference index of +/-0.43 (Fig. 3D). Neurons whose responses did not meet this criterion were classified as having intermediate preference (i.e., they tended to respond to *both* song modes) (Fig. 3A). Surprisingly, song mode preferences across neurons fell along a continuum (Fig. 3D), with 31% pulse-preferring, 31% sine-preferring, and the remaining 38% with intermediate preference. We re-analyzed 19,036 ROIs (regions of interest) spanning the central brain of 33 flies imaged pan-neuronally (Pacheco et al., 2020) according to the criteria established here, and found a similar, overlapping continuum (Supp. Fig 5C). We therefore conclude that the song mode preferences of the 28 WED/VLP cell types recorded here are representative of those of the entire auditory pathway recorded via whole brain imaging techniques.

### Auditory response variability varies by cell type

Within a cell class, the temporal dynamics of responses to song could vary across flies, even if the categorical preference for pulse vs. sine was consistent (e.g., Fig. 1G). To quantify this variability, we clustered the trial-averaged temporal traces to the three main types of stimuli presented, similar to a recent study (Pacheco et al., 2020), and agnostic to the cell type the recording came from. In other words, each fly recording was treated as a separate data point. Trial-averaged responses from WED/VLP neurons could be grouped into 13 clusters (Fig. 4A, see Methods), based on differences in the relative strength of excitation or inhibition, and the strength of adaptation.

**Figure 4.**
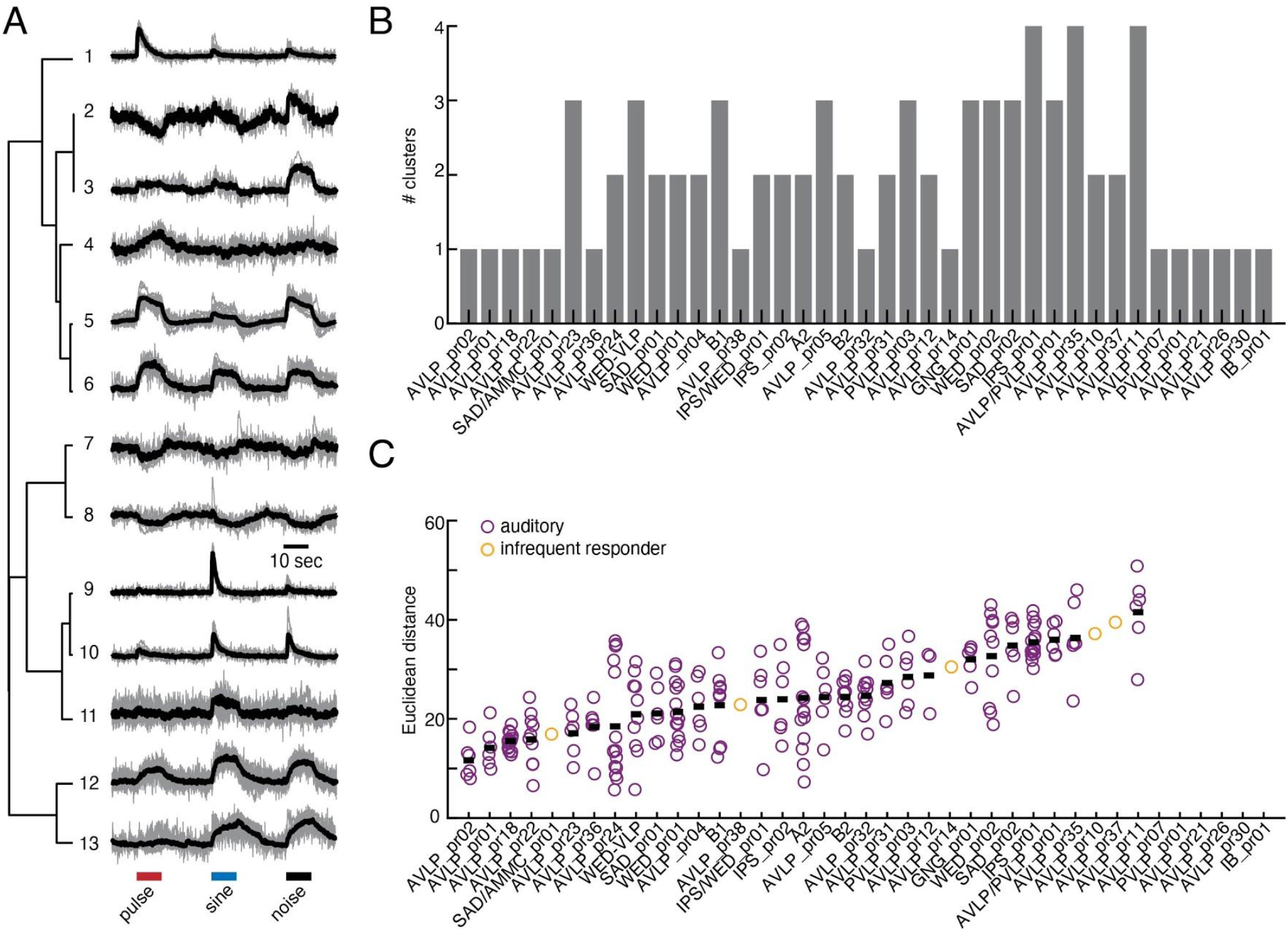
Across-animal variability of WED/VLP auditory responses. A) 13 different response types based on hierarchical clustering of calcium responses to pulse, sine, and noise stimuli. Trial-averaged responses from individual flies are shown in grey, and the mean within a cluster is shown in black. Each cluster contains the responses from 3-21 flies. B) The number of clusters into which each WED/VLP cell type’s responses were sorted. C) The Euclidean distances between every pair of responses within a WED/VLP cell class. Each dot represents the Euclidean distance between one pair, and black bars represent the mean within a cell class. We recorded only one response from each of the last six cell types on the x-axis.

We then determined how often responses from a given cell class (across flies) were grouped into the same cluster (Fig. 4B). The across-individual responses of 7 cell classes (AVLP_pr02, AVLP_pr01, AVLP_pr18, AVLP_pr22, AVLP_pr36, AVLP_pr32) were similar enough to each other to always be assigned to the same functional clusters, whereas the responses of 3 cell classes (IPS_pr01, AVLP_pr35, AVLP_pr11) were different enough to never be assigned to the same clusters. We further quantified inter-individual variability by calculating the Euclidean distance between responses from every pair of flies within each cell class (Fig. 4C), and found that, as expected, recordings from neurons that tended to fall into multiple clusters showed greater pairwise distances (Spearman’s rank correlation, *rho*=0.58, *p*<0.001), even though we controlled for developmental age, sex, receptivity state, and rearing conditions (see Methods). These results suggest that auditory responses from some cell types vary across individuals, with important implications for the processing of courtship songs among a population of flies (see Discussion).

### Variation in sine frequency and pulse rate tuning among WED/VLP neurons

We next examined sine frequency and pulse rate (or interpulse interval) tuning across our dataset (Fig. 3B, 5A,E). Behavioral data demonstrate that both males and females change their locomotor speed most in response to conspecific sine frequencies (across a wide range from 150-400 Hz) and pulse rates (peaked at 35-40 ms interpulse intervals) (Deutsch et al., 2019). However, among the WED/VLP population, many cell types were tuned to features outside of this conspecific song range (Fig. 5B,F). We analyzed sine frequency and pulse rate tuning for each fly, as responses can vary across individuals (Fig. 4), but we sorted tuning curves according to the overall selectivity of each cell type for sine versus pulse stimuli (Fig. 3D). For sine frequency, the majority of responses showed low-pass or band-pass tuning, with roughly equal numbers of responses in these two categories (Fig. 5C-D). Recordings from neurons we had characterized as pulse-preferring (Fig. 3D) predominantly showed band-pass frequency tuning centered on conspecific pulse carrier frequencies (e.g. around 250 Hz; (Clemens et al., 2018a)), whereas recordings from neurons characterized as sine-preferring or with intermediate preference showed either low-or band-pass tuning, with individual tuning curves tiling the broad range of frequencies present in conspecific sine song (100-400 Hz) (Arthur et al., 2013).

**Figure 5.**
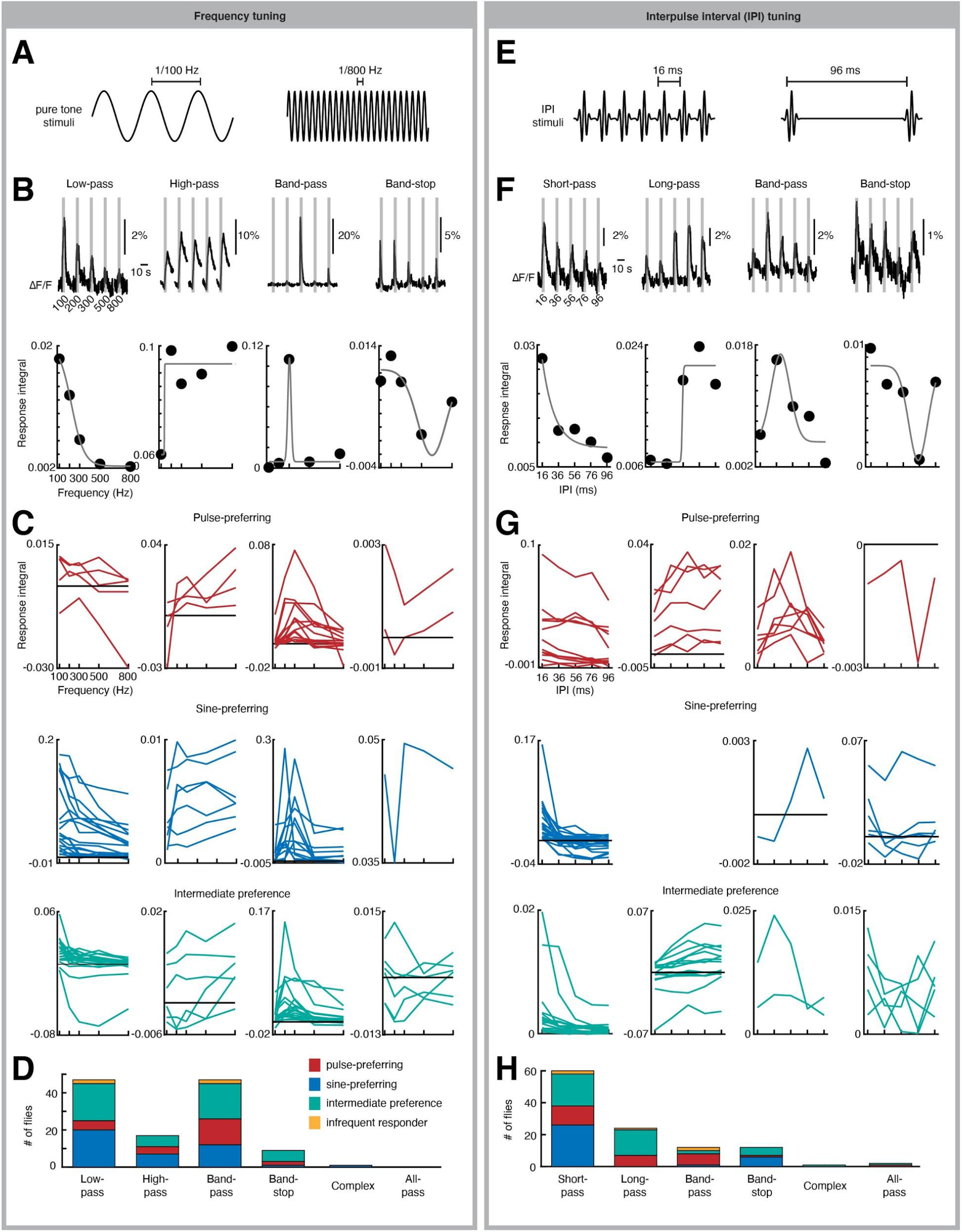
Frequency and pulse rate (interpulse interval) tuning. A) Example pure tone stimuli at the lowest and highest frequencies tested. B) Example recordings of responses to sine frequency stimuli. These individual responses (from four different flies) show high-pass, low-pass, band-pass, or band-stop tuning. Grey bars indicate when each stimulus was presented. The tuning curves are formed by measuring the integral of the responses to each frequency. C) Tuning curves for each fly recorded in the data set. Tuning curves are colored according to the song mode preference of each WED/VLP neuron type. D) Histogram of the frequency of tuning types across the dataset. Responses that were roughly equal for every stimulus were classified as all-pass, and responses that did not fit any other category were classified as complex (see Methods). E-H) Same as A-D for interpulse interval stimuli.

For pulse rate tuning, the majority of neurons exhibited short-pass responses (Fig. 5G-H). Whereas sine-preferring neurons were most likely to exhibit short-pass responses (consistent with these neurons preferring sinusoids over pulses since the shortest pulse rates begin to approximate sinusoids), pulse-preferring and intermediate-preference neurons showed much more diversity in pulse rate tuning. Surprisingly, relatively few recordings (2 flies from cell type AVLP_pr36 and 1 fly each from cell types SAD_pr01, SAD/AMMC_pr01, and AVLP_pr35) exhibited band-pass tuning for the conspecific pulse interval (35-40 ms). This is in contrast with cell types tuned for conspecific intervals found in brain regions downstream from the WED/VLP (e.g., pC2l (Deutsch et al., 2019) and vpoEN (Wang et al., 2020b)). Our results suggest that WED/VLP neuron responses may serve as a set of building blocks for generating band-pass tuning downstream in the pathway. We examine this hypothesis below by mapping connectivity between WED/VLP cell types and such downstream pulse song-tuned neurons.

### Spatial segregation of WED/VLP neurons by functional class within the VLP but not in projections to other brain areas

We next determined whether auditory neurons segregate by song mode preference either within the WED/VLP or in downstream target areas. In the AVLP, we found that pulse-preferring neurons most frequently targeted medial and anterior regions, whereas sine-preferring neurons were biased towards a posterior tract extending from ventral to dorsal areas (yellow in Fig. 6A). Neurons with intermediate preference targeted areas innervated by both pulse- and sine-preferring lines, whereas non-auditory lines most frequently targeted the lateral half of the AVLP. In the PVLP, projections of sine-preferring neurons were most common in an anterior region, roughly in the middle of the medial-lateral and dorsal-ventral axes, whereas projections of pulse-preferring neurons were found more medially and dorsally (yellow in Fig. 6B). In contrast to the AVLP, intermediate-preference lines innervated a region separate from the major areas innervated by pulse- and sine-preferring neurons. Non-auditory lines tended to innervate more lateral PVLP regions than auditory lines (yellow in Fig. 6B). In contrast to VLP projections, WED projections showed less pronounced differences between pulse-vs. sine-preferring, and auditory vs. non-auditory response categories (Fig. 6C). Taken together, these results suggest that sine and pulse preference largely arises in the VLP in separate anatomical regions, and that this transformation in stimulus preference likely occurs between WED projection and VLP projection neurons.

**Figure 6.**
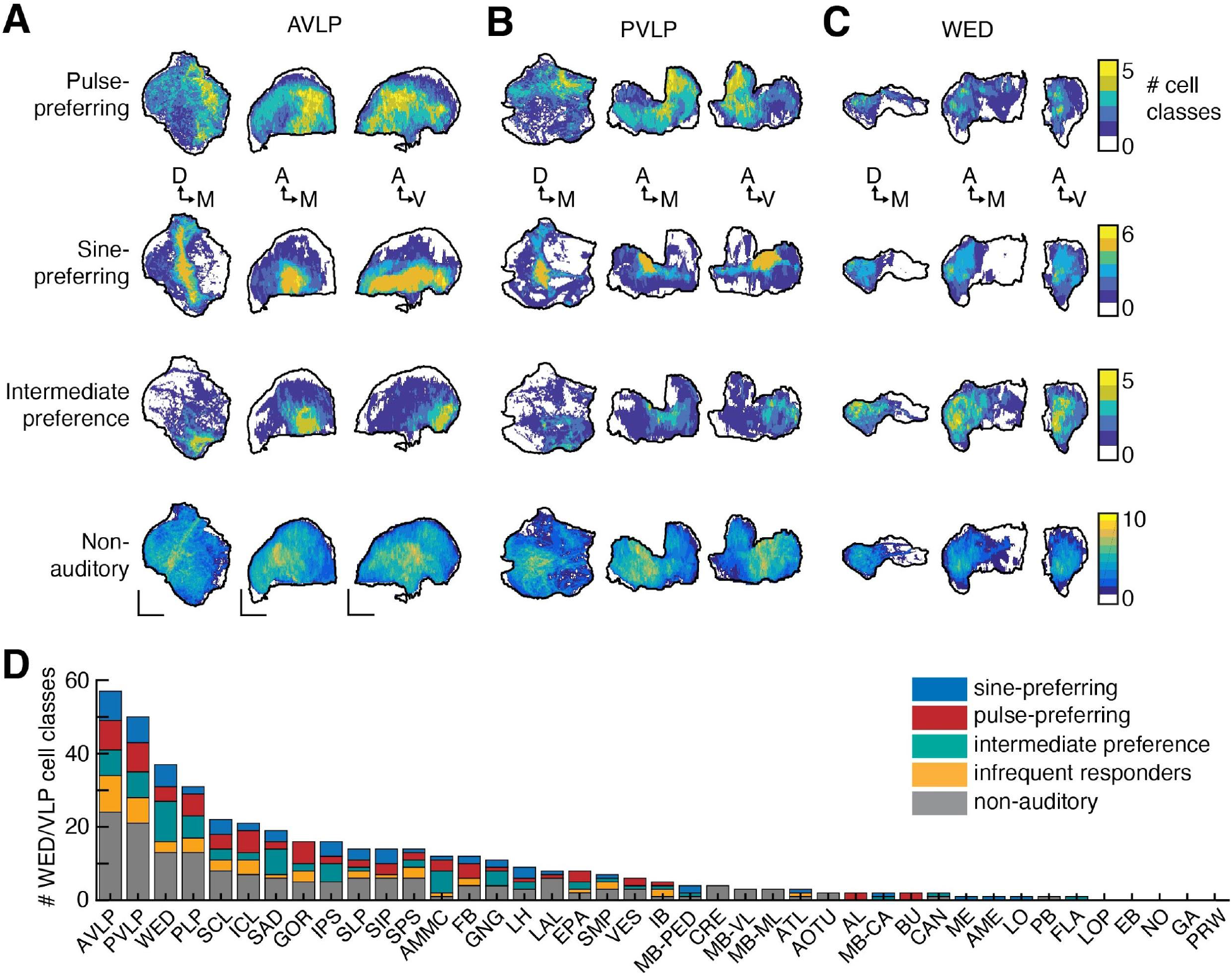
Auditory neurons are spatially segregated in the VLP according to song mode preference, but project to a diversity of brain areas. A-C) 2D projections of the maximum number of WED/VLP cell classes with expression in voxels within the right hemisphere of the AVLP (A), PVLP (B), and WED (C) neuropils. Scale bars = 25 microns. D) Histogram of the total number of WED/VLP cell classes with innervation in each neuropil. Neuropil abbreviations are listed in Table 1.

In contrast with the spatial segregation of functional class found in the VLP, WED/VLP auditory neurons sent projections throughout the central brain, independent of song mode preference (Fig. 6D; Supp. Fig. 6), including to regions considered primarily visual (ie, posterior lateral protocerebrum or PLP) and olfactory (i.e., lateral horn (LH)). Auditory neurons also frequently targeted neuropils known to be rich in the processes of sexually dimorphic neurons, such as the superior clamp (SCL), inferior clamp (ICL), and inferior posterior slope (IPS). These findings are consistent with results of pan-neuronal imaging that found auditory responses throughout 33/36 neuropils of the central brain (Pacheco et al., 2020). Ten of 12 WED/VLP cell classes with projections in the AMMC responded to auditory stimuli in our screen (Fig. 6D), which is consistent with the AMMC being the primary target of auditory receptor neurons (Matsuo et al., 2016). For other neuropils, there were no obvious differences in innervation patterns based on auditory response type or lack thereof (Fig. 6D, Supp. Fig. 6), suggesting auditory activity cannot be predicted based solely on which brain regions a WED/VLP neuron targets.

A portion of the PVLP and posterior lateral protocerebrum (PLP) innervated by many of our WED/VLP neurons contains the optic glomeruli (Otsuna and Ito, 2006; Panser et al., 2016; Strausfeld and Gronenberg, 1990; Wu et al., 2016). Neurons innervating the optic glomeruli play important roles in a number of visual behaviors. We found that 11 auditory WED/VLP neurons and 6 infrequently responding WED/VLP neurons contained processes in at least one optic glomerulus (Supp. Fig. 7; see Methods). The two optic glomeruli with the most innervation by our WED/VLP cell classes were LC9 and LC11, both containing small object-detecting visual neurons that might be important for courtship behaviors (Bidaye et al., 2020; Keleş and Frye, 2017; McKinney and Ben-Shahar, 2019). The LC9 glomerulus is exclusively innervated by neurons that prefer pulse stimuli whereas the LC11 glomerulus is mostly innervated by neurons that prefer sine stimuli (Supp. Fig. 7). As sine song is produced at close range to females while pulse is produced farther away (Coen et al., 2014), it is interesting to note that LC11 neurons are tuned for small objects (Keleş and Frye, 2017) while LC9 neurons provide inputs to P9 neurons that drive female-directed turning and walking in courting males (Bidaye et al., 2020). However, since this analysis is based on LM images, it remains to be determined whether auditory WED/VLP neurons send outputs to or receive inputs in optic glomeruli.

### The connectome of *Drosophila* auditory neurons

We uncovered a continuum of song mode preference strengths across the population of WED/VLP auditory neurons (Fig. 7A). Two possible models of network organization would be consistent with such a continuum. In the first model, pulse- and sine-preferring neurons constitute separate pathways with their own intermediate-preference neurons, and hierarchical organization within each pathway sharpens tuning to generate strong song mode preference. Support for model 1 comes from work on higher-order pulse song preferring neurons ((Deutsch et al., 2019; Wang et al., 2020a, 2020b)). Two subsets of neurons (pC2l and vpoEN) were found to be strongly pulse preferring, and to connect with descending neurons (neurons with dendrites in the brain and axon terminals in the ventral nerve cord) that drive sex-specific behaviors. This suggests song mode preference is built up within a pathway that separates sine and pulse processing into distinct streams to drive disparate behaviors. In the second model, the continuum of responses arises through extensive interconnectivity between cell types with different preferences. In model 2, inhibitory interactions between cell types with different song mode preferences could act to sharpen tuning. We tested which network organization underlies auditory processing in the *Drosophila* brain by mapping synaptic connectivity among all auditory neurons (both WED/VLP neurons studied here and auditory neurons characterized in prior studies). We identified and proofread these neurons in *both hemispheres* of a whole brain electron microscopic volume (FlyWire) (Dorkenwald et al., 2020; Zheng et al., 2018) (Fig. 2, 7B), determined the number of synaptic connections via automated synapse detection (Buhmann et al., 2019)) (Supp. Fig. 8), and used a new deep learning-based classifier to predict the neurotransmitter used by each cell type (Eckstein et al., 2020) (Supp. Fig. 9). To relate connectivity with function, we labeled each cell type according to its song mode preference either defined in this study (Fig. 3D) or in prior work (Deutsch et al., 2019; Wang et al., 2020a, 2020b), bearing in mind that there is diversity in preference strength across the population (Fig. 3D). This revealed extensive interconnectivity between auditory cell types with different song mode preferences (Fig. 7C), more consistent with model 2.

**Figure 7.**
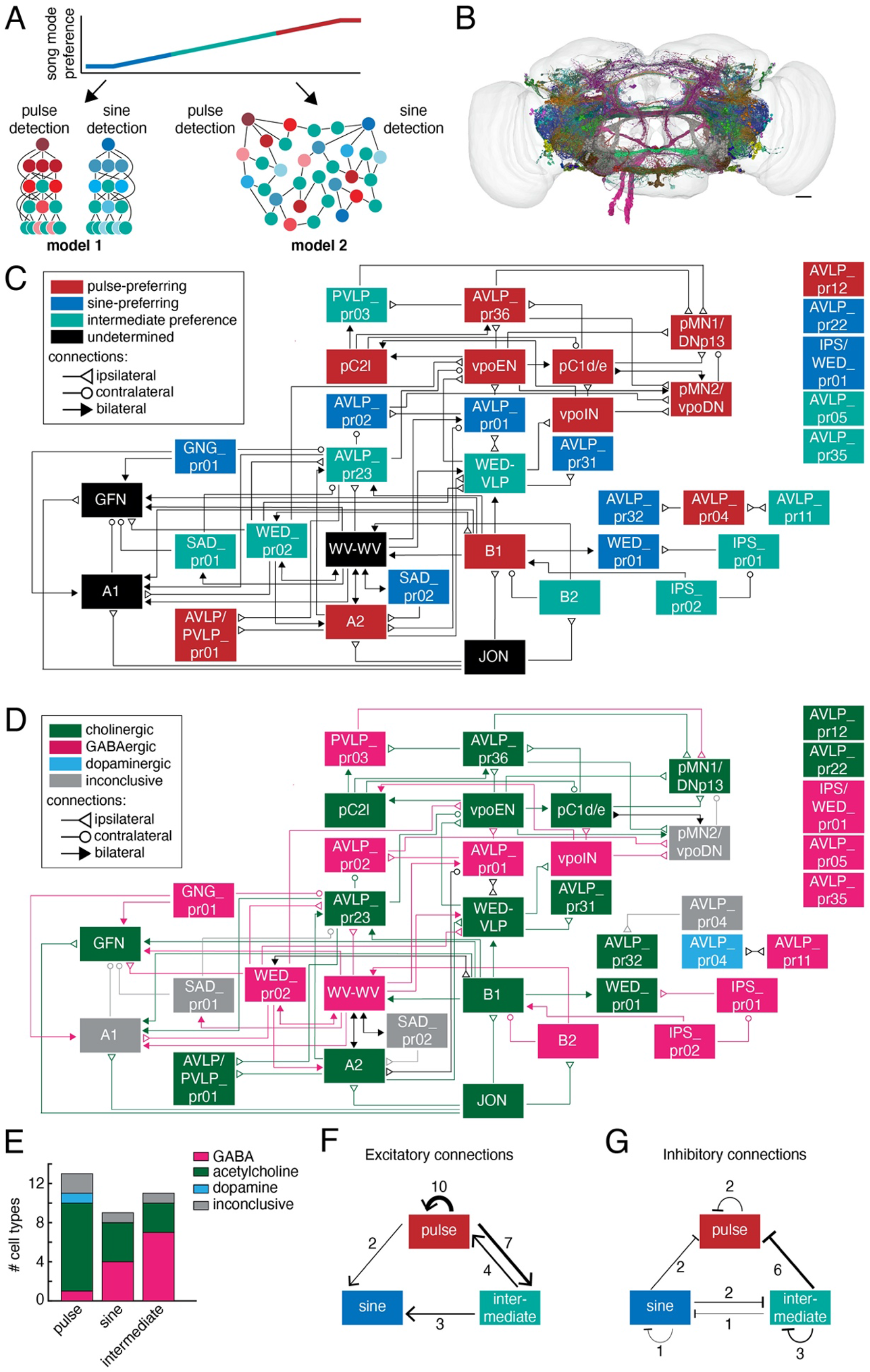
The auditory connectome. A) Two models for the organization of auditory pathways underlying the observed continuum of song mode preferences. Each circle represents a cell type, and each line represents synaptic connections. Red = pulse-preferring, blue = sine-preferring, and green = intermediate preference (see Fig. 3D). The shading of red and blue neurons indicates the strength of pulse or sine preference, respectively. In model 1, neurons selective for pulse and sine are separated into distinct pathways, with neurons of intermediate preferences playing roles in both pathways (left). Within-mode connections sharpen tuning for song features. In model 2, neurons of different song mode preferences are highly interconnected at all levels without hierarchical organization (right). Downstream neurons may pool the responses of diversely tuned neurons to establish selectivity for a variety of song parameters. B) To test these models, we examined synaptic connections among auditory neurons (N=496 neurons from 35 cell types) in both hemispheres of an EM volume of an entire female fly brain. Scale bar = 25um. C) Schematic of the synaptic connectivity of auditory neurons. Each box represents a cell type, and each line represents a synaptic connection (see Methods for connection criteria). The song mode preference of A2, B1, B2, and WED-VLP come from recordings from the split-GAL4 lines labeling those neurons (Supp. Fig. 5B; Table 2), and the song preferences of several higher-order and descending neurons (pC2l, vpoEN, vpoIN, pC1d/e, pMN1/DNp13, and pMN2/vpoDN) come from previous studies (Zhou et al., 2014; Deutsch et al., 2019; Wang et al., 2020a, 2020b). Cell types shown in black boxes have not had their sine/pulse preference determined. Five cell types formed no connections with other neurons in the dataset. See Methods for how the connectivity diagram was formed from identified reconstructions in FlyWire.ai. In contrast to results using the hemibrain EM volume (Wang et al., 2020a), we did not find strong connections between pC2l and pMN1/DNp13. Instead, pC2l neurons provided inputs with fewer than 10 synapses per neuron onto pMN1/DNp13 in FlyWire (Supp. Fig. 8). However, our FlyWire results replicate other findings based on hemibrain data, including strong synaptic inputs from vpoEN and vpoIN onto pMN2/vpoDN (Wang et al. 2020b). D) Same schematic as in (C) but with cell types and synaptic connections colored by neurotransmitter (Supp. Fig. 9C). Bidirectional connections between cells with different neurotransmitters are shown in black. Since the neurotransmitter results for AVLP_pr04 suggested two possible cell types, and since synaptic connectivity was different for these two cell types, AVLP_pr04 is split into a dopaminergic cell type and a cell type with inconclusive neurotransmitter. E) Histogram of the number of cell types with each song mode preference and neurotransmitter. F-G) Summaries of excitatory (F) and inhibitory (G) connections among auditory neurons. Line thickness represents the relative number of connections.

The auditory pathway begins in the Johnston’s Organ (JO) of the antenna, where JO-A and JO-B neurons detect sounds and relay information to the AMMC (Kamikouchi et al., 2009). Information then splits in the AMMC, into a pathway including the giant fiber neuron (GFN) and A1, which receive inputs largely from JO-A neurons, integrate this information with other sensory streams, and drive escape maneuvers ((Allen et al., 2006; von Reyn et al., 2014), and another pathway including neurons (A2, B1, and B2) that receive inputs from a mix of JO-A and JO-B neurons and are considered to constitute the first stages of the courtship song processing pathway (Kim et al., 2020; Vaughan et al., 2014; Yamada et al., 2018). We found that two of these cell types shows strong song mode preference (A2 and B1; Fig. 3D), suggesting that song mode preference arises at the first relay of the pathway and not via sharpening of tuning along a hierarchical network, inconsistent with model 1. In addition, building on recent connectomic work (Dorkenwald et al., 2020), we found a number of connections between the song and escape pathways. These connections are in one direction (from the song pathway to the escape pathway) and are largely inhibitory (e.g., GNG_pr01, WED_pr02, and WV-WV inhibit the GFN, while a specific subset of B1 neurons excites the GFN (Supp. Fig. 10)).

We also found little evidence of hierarchical organization of the network at a purely structural level. Using the binarized neural connectivity matrix (Supp. Fig. 8), we computed three measures of network hierarchy: orderability, feedforwardness, and treeness (Supp. Fig 11A). These metrics evaluate the fraction of neurons belonging to network cycles (or recurrent loops in the network), the overall contribution of cycles to paths through the network, and the extent to which the network diverges (fans out) from a set of root nodes, respectively (Corominas-Murtra et al., 2013). While the network could be condensed into a feedforward network of connected components (Supp. Fig. 11B), with approximately half of the neurons uniquely orderable (Supp. Fig. 11C), paths through the network were nonetheless dominated by recurrent connections, and the network tended to neither fan out nor converge from a set of root neurons. The network deviated only marginally in each hierarchy metric from random networks with matched degree distributions (Supp. Fig. 11C-E), with values markedly less than those of a perfectly feedforward, tree-like network. Thus, the network connectivity structure, even when not considering the song mode preference of individual cell types, appears to exhibit little hierarchical organization beyond what one would expect from chance.

We found many examples of connections between cell types with distinct song mode preferences (Fig. 7C-D). For example, intermediate preference WED-VLP neurons receive inputs from all preference classes, and in turn send excitatory projections to two pulse-preferring (vpoEN and vpoIN) and two sine-preferring (AVLP_pr01 and AVLP_pr31) neurons. The pulse-preferring B1 neurons send outputs to and vpoEN receives inputs from at least one cell type with intermediate and sine preferences, but no pulse-preferring neurons. The WED_pr01 cell type, which is strongly sine-tuned, receives excitatory projections from pulse-tuned B1 neurons, and inhibitory projections from intermediate preference IPS_pr01 neurons. Finally, cell types with both intermediate preference (WED-VLP, WED_pr02, and AVLP_pr23) and sine preference (AVLP_pr01, AVLP_pr02) provide input to the sub-network of higher-order pulse-preferring neurons (vpoEN, vpoIN, pC1d, pC2l, pMN1/DNp13, pMN2/vpoDN, and AVLP_pr36). While we acknowledge that the auditory connectome generated here is incomplete, these data nonetheless reveal a lack of segregation of neurons into pulse and sine song processing pathways (consistent with model 2), and uncover surprising transformations between inputs and outputs to generate a wide diversity of tunings.

We also observed a mix of excitatory and inhibitory connections throughout the network (Fig. 7D). Overall, pulse-preferring neurons were overwhelmingly cholinergic, whereas sine-preferring and intermediate preference neurons included both cholinergic and GABAergic cell types (Fig. 7E). We found one dopaminergic neuron in the pulse-preferring AVLP_pr04 cell type. Inhibitory connections occurred between almost every possible pair of preference classes (Fig. 7G), indicating that inhibition may play multiple roles in shaping song tuning. Inhibitory connections were most common between intermediate preference neurons and pulse-preferring neurons (Fig. 7G), suggesting that this connection type may be especially important for establishing responses to pulse song features. Sine-preferring neurons inhibited both pulse and intermediate preference neurons, suggesting one major role for sine-preferring inhibitory neurons is to oppose responses to sine song (Fig. 7G).

The most frequent connection type was pulse-preferring neurons exciting other pulse-preferring neurons (Fig. 7F) - many of these connections were in the sub-network of pulse-preferring neurons that includes descending neurons pMN1/DNp13 and pMN2/vpoDN (Mezzera et al., 2020; Wang et al., 2020b). Pulse-preferring neurons also excited intermediate-preference neurons (Fig. 7F), while intermediate-preference neurons excited both pulse- and sine-preferring neurons, raising the question of the role of intermediate auditory preferences in shaping downstream tuning (Fig. 7F). Although roughly half of the sine-preferring cell types were excitatory, we did not identify any of their downstream targets in our auditory dataset (Fig. 7E-F). Nonetheless, this reveals that there are excitatory pathways for sine processing, with downstream, descending targets yet to be identified.

Taken together, our functional and anatomical results support a novel network organization for auditory processing: synaptic connections (both excitatory and inhibitory) between cell types with different tunings contribute to generating a continuum of song mode preferences (Fig. 3D). Although the overall network organization is non-hierarchical (Supp. Fig. 11), when considering song mode tuning, we find a clear sub-network of pulse-preferring neurons. Therefore even though the neural network organization of the auditory pathway is more consistent with model 2, we also find some evidence in support of separate nodes for decoding pulse versus sine song.

## DISCUSSION

Here we have identified and characterized 24 new central auditory neuron types in the *Drosophila* brain. Prior work had focused on either early or higher-order auditory neurons, leaving open the characterization of intermediate cell types in the auditory pathway, and no studies had systematically characterized tuning for the two modes of courtship song. By additionally examining synaptic connectivity among all known auditory neurons using a new connectomic resource, FlyWire, we discovered extensive interconnectivity between cell classes with different song mode preferences. These results rule out a model in which the two main song modes are processed in separate pathways, and instead support a model in which interconnectivity contributes to generating a continuum of song mode preferences. Such a continuum has been observed in the sensory systems of fish (Sproule and Chacron, 2017; Thompson and Scott, 2016) and sensorimotor cortices of rodents ((Minderer et al., 2019); (Raposo et al., 2014); (Saleem et al., 2013)), and thought to support encoding of a larger array of variables than would be possible with categorical representations (in other words, a more efficient representation). In the fly auditory system, neurons with intermediate preference could encode sound intensity, location, and long timescale features, such as the variable sequence of pulses and sines that make up song bouts (Coen et al., 2014) - behavioral studies demonstrate that females are sensitive to song bout features such as the average duration of song bouts over timescales of minutes (Clemens et al., 2015). The continuum of song preferences we discovered among WED/VLP neurons matches the continuum found throughout the entire brain (Supp. Fig. 5), suggesting that these neurons are representative of the larger auditory population, including those neurons that drive song-responsive behaviors. Many descending neurons that control locomotion have dendrites in a number of neuropils with auditory activity, including WED/VLP (Namiki et al., 2018; Rayshubskiy et al., 2020). Sampling from a continuum of responses to sine and pulse stimuli would provide these DNs with the ability to respond to a wide array of song patterns.

We identified new auditory neurons by targeting specific cell classes within the WED/VLP, putatively downstream from previously known auditory neurons (Lai et al., 2012; Matsuo et al., 2016). In the auditory connectome we built using FlyWire (Dorkenwald et al., 2020), which allows proofreading of EM reconstructions in an entire female brain (Zheng et al., 2018), we were able to examine inter-hemispheric connections, as well as ipsilateral vs. contralateral connection patterns (Fig. 7C-D). The circuit map provides hypotheses for how flies localize sounds in space ((Morley et al., 2012); (Batchelor and Wilson, 2019)). However, several cell types formed no synaptic connections with other auditory neurons in our data set, and many other neurons had synaptic inputs but no outputs (and vice versa). We also found one pathway, including AVLP_pr32, AVLP_pr04, and AVLP_pr11, that did not synaptically connect with any other auditory neurons. These results indicate that despite our attempts at comprehensively identifying new auditory WED/VLP neurons, the auditory connectome is still incomplete. In support of this, we were not able to identify split-GAL4 lines for several auditory VLP neurons characterized previously (Clemens et al., 2015). How many neurons may be still unknown in the auditory pathway is difficult to estimate. Continued circuit tracing in EM combined with functional imaging of sparse and specific driver lines, as we have done here, should ultimately fill out the auditory connectome. It is important to note that since song representations are present in 33 out of 36 central brain neuropils (Pacheco et al., 2020), we expect auditory circuits to be highly interconnected to circuits of other modalities and functions. Indeed, the new auditory neurons we discovered send projections throughout the brain, including to olfactory, visual, and pre-motor areas, suggesting a role in multisensory integration.

We found that the responses of some auditory cell types were highly stereotyped across animals, whereas others were more variable. Pan-neuronal imaging also found across-animal variability in auditory responses (Pacheco et al., 2020) but could not map this variability to individual cell types, as we have done here. In the mushroom body, Kenyon cells (KCs) of a single type also have variable odor responses across animals (Murthy et al., 2008) which arises through stochasticity in connectivity between antennal lobe projection neurons and KCs (Caron et al., 2013; Zheng et al., 2020)). This stochasticity might be important for generating variation in odor preferences or odor learning capacity across animals. Across-animal wiring differences in the central complex lead to differences in left vs. right turn biases (Buchanan et al., 2015), which may be important for generating diversity in navigation strategies across a population of flies. The role of variability in auditory responses is currently unknown, but it may function to generate variation in song preference across animals, potentially useful for diversifying female mate selection preferences.

Our study uncovered differences in frequency and pulse rate tuning between sine- and pulse-preferring neurons. Sine-preferring neurons exhibited diverse frequency tuning but more uniform rate tuning (Figs. 5D,H). The short-pass rate tuning in sine-preferring neurons is consistent with their preference for sinusoidal, or continuous, stimuli. In contrast, pulse-preferring neurons exhibited more uniform frequency tuning but diverse rate tuning (Figs. 5D,H). Pan-neuronal imaging of neurons in the AMMC and WED neuropils found tonotopy in JON axons that was maintained in the AMMC and potentially sharpened in the WED (Patella and Wilson, 2018). However, the extent to which the sine frequency and pulse rate tuning of WED/VLP neuron types is inherited from inputs and/or shaped downstream of the JONs remains unknown. Very few WED/VLP neurons showed band-pass tuning for conspecific pulse rate, suggesting that downstream processing is required for generating such selectivity, as is observed in neurons such as pC2l or vpoEN (Deutsch et al., 2019; Wang et al., 2020b; Zhou et al., 2015). This pattern would be consistent with that of vertebrate auditory systems, in which frequency tuning arises in the periphery, whereas tuning for temporal sound features, such as pulse rate, arises through central computations (Winer and Schreiner, 2005). An advantage of computing conspecific pulse rate centrally could be the flexibility to rapidly adapt auditory preferences to evolutionary change in song parameters, particularly following speciation events.

Mapping synaptic connectivity among auditory neurons revealed a high degree of intermixing between preferences for the two song modes (Fig. 7C). One such example is inhibitory interactions that refine tuning. For instance, the convergence of sine-preferring inhibitory inputs and excitatory inputs with intermediate song mode preference would result in stronger pulse-preference than the excitatory inputs alone. The use of inhibition to sharpen central auditory tuning is widespread across taxa. In grasshoppers and katydids, the convergence of narrowly-tuned inhibition with broadly-tuned excitation shapes frequency tuning (Romer et al., 1981; Stumpner, 2002). In mammalian auditory circuits, sideband inhibition, in which stimuli just outside of a neuron’s preferred range elicit inhibition, also sharpens frequency tuning in central neurons (Woolley and Portfors, 2013). We found inhibitory connections between almost every possible combination of song-mode preference classes (Fig. 7G), suggesting that inhibition may play a number of roles in shaping song tuning in flies.

In conclusion, prior work on auditory systems has led to ideas about segregation of processing into separate streams for different sound categories (Rose and Brenowitz, 2002; Wyttenbach et al., 1996), but these ideas do not address how animals might encode patterns or sequences of sound elements. Further, while separate streams may be useful for detecting different classes of stereotyped acoustic signals, such segregation may not be sufficient for analyzing rapidly varying sequences of elements, as occur in fly and some types of bird songs (Coen et al., 2014; Cohen et al., 2020). In these cases, encoding longer timescale patterns, such as the recent history of interleaved song modes, may benefit from drawing upon neural responses with a range of song mode preferences. Future work should examine the ways in which this network is read out by downstream neurons to drive acoustic communication behaviors.

## ACKNOWLEDGEMENTS

We thank Xiao-Juan Guan for help with crosses during split-GAL4 generation; David Deutsch for help with calcium imaging pre-analysis and for identifying pC2l neurons in FlyWire; Sebastian Seung for assistance with FlyWire infrastructure; Greg Jefferis for feedback on the cell type naming scheme and Alex Bates and Greg Jefferis for helpful discussions and ideas with regard to neurotransmitter analysis; Nat Tabris for help with connectivity analysis; Lucas Encarnacion-Rivera, Ben Silverman, Jay Gager, Merlin Moore, Selden Koolman, James Hebditch, Sarah Morejohn, Kyle Willie, Austin Burke, Celia David, and Szi-chieh Yu for proofreading FlyWire neurons; Marta Costa, Jan Clemens, and Istvan Taisz for comments on the manuscript; and the Janelia Fly Core, FlyLight, and Scientific Computing groups. A.N. thanks Gerry Rubin, in whose lab he performed this work, for his support and encouragement.

## AUTHOR CONTRIBUTIONS

CAB, CM, and MM designed the study; CM, CAB and AN screened for split-GAL4 lines in collaboration with BJD; CAB collected and analyzed split-GAL4 neural recording data, with assistance from DP; DP collected and analyzed pan-neuronal imaging data and performed neuropil registrations; CAB conducted connectome analyses with assistance from SD; RP performed analysis on the hierarchical structure of the auditory connectome; NE and JF performed neurotransmitter analysis; CAB and MM wrote the manuscript, with feedback from all authors.

## METHODS

### Fly stocks

The split-GAL4 system (Luan et al., 2006; Pfeiffer et al., 2010) was used to express the activation domain (AD) and the DNA-binding domain (DBD) of GAL4 under the separate control of two genomic enhancers, to obtain the intersection of their expression patterns, with the goal of obtaining a sparser pattern. Split-GAL4 stocks were gifts of Gerald Rubin and Barry Dickson (Dionne et al., 2018; Pfeiffer et al., 2010; Tirian and Dickson, 2017). 20xUAS-GCaMP6s, td-Tomato/CyO was generated by Diego Pacheco (Pacheco et al., 2020). The genotype of imaged flies was 20x-UAS-GCaMP6s, td-Tomato/GAL4-AD; GAL4-DBD/+, with the “+” originating from NM91 stocks. For expression pattern staining, split-GAL4s were combined with 20xUAS-CsChrimson-mVenus trafficked in attP18, except that 6 lines (17963, 21914, 23281, 23627, 28822, 29146) were shared from a different project that instead used pJFRC51-3XUAS-IVS-syt::smHA in su(Hw)attP1,pJFRC225-5XUAS-IVS-myr::smFLAG in VK00005, and one (27932) used pJFRC200-10XUAS-IVS-myr::smGFP-HA in attP18, pJFRC216-13XLexAop2-IVS-myr::smGFP-V5 in su(Hw)attP8. For multicolor flip-out (MCFO) staining, all were crossed to MCFO-1 [pBPhsFlp2::PEST (attP3);; pJFRC201-10XUAS-FRT>STOP>FRT-myr::smGFP-HA (VK0005), pJFRC240-10XUAS-FRT> STOP>FRT-myr::smGFP-V5-THS-10XUASFRT>STOP>FRT-myr::smGFP-FLAG (su(Hw)attP1)]. Flies were kept at 25C with a 12h:12h light:dark cycle.

### Split-GAL4 creation

GAL4 images from the Rubin and Dickson collections (Jenett et al., 2012; Pfeiffer et al., 2010; Tirian and Dickson, 2017) were visually screened for lines targeting neurons in the WED and AVLP. For each cell type, a color depth MIP mask search (Otsuna et al., 2018) was conducted to find other GAL4 lines with expression in similar cells. Split-GAL4 hemidrivers for these lines were crossed in various combinations to find a combination that targeted the cell type of interest but with sparse expression elsewhere, determined by expression and staining of mVenus in one female fly of genotype 20xUAS-csChrimson::mVenus (attP18)/w; Enhancer-p65ADZp (attP40)/+; Enhancer-ZpGAL4DBD (attP2)/+. 65 combinations were chosen by this method, representing over 50 cell types. In some cases, lines targeting neurons with similar gross morphology revealed one line with a commissure (see SS16374 and SS27885; SS41728 and SS41730 in Supp. Fig. 1). For these examples, we characterized both lines due to the difficulty in determining whether they were truly similar cell types. Split-GAL4 combinations were then double balanced and combined in the same fly strain to make a stable split-GAL4, for which expression pattern staining was carried out in an additional female to confirm the expression seen in the initial screen. For multiple split-GAL4 lines targeting morphologically similar neurons, one (or sometimes a few) were selected according to the following criteria: least off-target expression in the central brain, largest number of neurons with expression, and/or most reliable calcium responses to acoustic stimuli across flies. Expression was further confirmed in multiple flies by multicolor flip-out staining (see below). All lines generated here and the corresponding image stacks are available via https://splitgal4.janelia.org.

### CNS immunohistochemistry & imaging

Brains and ventral nervous systems were dissected and stained using published methods (Aso et al., 2014; Nern et al., 2015; Wu et al., 2016). Antibodies used were rabbit anti-GFP (1:500, Invitrogen, #A11122), mouse anti-Bruchpilot (1:50, Developmental Studies Hybridoma Bank, University of Iowa, mAb nc82), Alexa Fluor 488-goat anti-rabbit (1:500, ThermoFisher A11034), and Alexa Fluor 568-goat anti-mouse (1:500, ThermoFisher A11031). Serial optical sections were obtained at 1µm intervals on a Zeiss 700 confocal with a Plan-Apochromat 20x/0.8NA objective. A detailed description of the staining and screening protocol is available at https://www.janelia.org/project-team/flylight/protocols under “IHC - Adult Split Screen.”

### Stochastic labeling

For multicolor flip-out (MCFO) stochastic labeling (Nern et al., 2015), approximately 8 females per split-GAL4 line received a 15 min heat shock at 37°C at 1-3 days old, and were dissected at 6-8 days. A detailed description of the MCFO protocol can be found at https://www.janelia.org/project-team/flylight/protocols under “IHC - MCFO.”

### Image processing

Images were adjusted for brightness and contrast without obscuring data. Images were processed in ImageJ (https://imagej.nih.gov/ij/) and Photoshop (Adobe Systems Inc.). Where noted, neurons were rendered and segmented from confocal stacks with VVDviewer software (https://github.com/takashi310/VVD_Viewer) (Wan et al., 2009, 2012) to visualize them in isolation. For this rendering and for computational alignment of brain images used where noted, brain images were registered using the Computational Morphometry Toolkit (Rohlfing, 2011) to a standard brain template (“JFRC2014”) that was mounted and imaged with the same conditions. Segmented image stacks are available at https://splitgal4.janelia.org.

### Neuropil innervation

We registered the neuropil (Ito et al., 2014) and optic glomeruli (Panser et al., 2016) maps to the unisex JRC2018 template by first bridging from IBNWB to JFRC2 (Bates et al., 2020), and then JFRC2 to unisex JRC2018 (Bogovic et al.). To determine which neuropils each neuron type targeted, we used the segmented neuron stacks with manually removed somata. We then calculated the percentage of the neuron’s voxels in each neuropil and optic glomerulus. In Fig. 6D and Supp. Fig. 3, 6, and 7 we report all neuropils containing at least 1% of each segmented neuron’s total volume.

For naming auditory neurons not previously described as auditory, we used the neuropil in which the segmented neuron had the greatest percentage of expression. If a neuron had nearly equal (<1% difference) expression in two neuropils, we used both neuropil names. A two digit number was added to the neuropil name in sequential order based on the split-GAL4 line number (Table 2). To disambiguate our names from those of the hemibrain project (Xu et al., 2020) (as our neuron types/cell classes can likely be further segregated into subtypes based on both morphological and connectivity differences within a line), we added a ‘pr’ before each number for Princeton.

### EM reconstructions

We identified individual neurons in the FAFB volume using FlyWire.ai (Dorkenwald et al., 2020). Neurons were proofread using the editing tools in FlyWire to add missing pieces of arbor and remove incorrect pieces, focusing on the main backbone of a neuron, not attempting to add very small missing twigs. Sister cells from the same cell type were used to verify that a cell’s overall morphology appeared generally correct. WED/VLP reconstructions were proofread by 12 proofreaders consisting of both scientists and expert tracers from the Seung and Murthy labs. To find all candidates belonging to a given cell type, we found an EM cross-section of the primary neurite tract, and investigated every cell in the cross-section. We compared candidate reconstructions with LM images of our auditory lines (Supp. Fig. 1-2). We were conservative with which reconstructions we included for each cell type; if a reconstruction did not match neurons present in MCFO images (Supp. Fig. 2), we excluded it. Since a limited number of MCFO images were available for each line, it is possible that we may have missed some EM reconstructions belonging to each line. Two auditory cell types (AVLP_pr18 and AVLP_pr24) did not have enough cell-type resolution in MCFO to enable individual EM reconstruction identification and were excluded from the synaptic connectivity mapping. EM reconstructions for pC1d/e, pMN1/DNp13, and pMN2/vpoDN came from a previous study (Deutsch et al., 2020). We identified vpoEN and vpoIN by inspecting the inputs to pMN2/vpoDN neurons (Wang et al., 2020b). pC2l reconstructions were identified by D. Deutsch (personal communication).

### Synaptic connectivity among auditory neurons

We used FAFB reconstructions corresponding to previously known auditory neurons (Dorkenwald et al. 2020), higher-order auditory neurons (Deutsch et al., 2020; Wang et al., 2020b), and all auditory WED/VLP neurons for which we found EM reconstructions. We then found all automatically detected synapses between every pair of individual neurons (Buhmann et al., 2019). We omitted synapses that may have been identified multiple times by eliminating synapses between a given pair for which the presynaptic site was within 150 um of another presynaptic site. We ignored synapses from one neuron onto itself and between individual neurons of the same cell type, due to the high rate of false positives in these cases (Dorkenwald et al., 2020). For the remaining indicated synaptic connections, we set a threshold of 15 synapses for a likely true connection between any pair of neurons. Applying our criteria to early auditory neurons results in the same connectivity results among the neurons in Dorkenwald et al. (2020), with one exception. Dorkenwald and colleagues found that one automatically indicated connection between cell types was due to a high number of inverted synapses (ie, automatically detected presynaptic partner was determined to be postsynaptic by a human observer). Our connectivity analysis method is unable to confirm synapse directionality, and it is not yet known how often this type of false positive occurs. Five WED/VLP cells (AVLP_pr05, AVLP_pr12, AVLP_pr22, IPS/WED_pr01, AVLP_pr35) did not have connections meeting these criteria with any other cell type in our dataset.

### Analysis of hierarchical network structure

We binarized the connection matrix between individual neuron pairs (Supp. Fig 8), keeping only connections where at least 15 synapses between the two neurons had been detected in the connectomic analysis, and then computed orderability, feedforwardness, and treeness of the resulting directed binary network *G*, as described in (Corominas-Murtra et al., 2013). We first computed the “node-weighted condensed graph,” or graph condensation, *G*_*C*_ (Supp. Fig. 11B) of the original network by identifying all strongly connected components {*v*_*i*_} of *G* (sets of neurons all mutually reachable from one another via at least one directed path) and retaining whether or not at least one directed connection existed between each pair of components. Thus, *G*_*C*_ is a directed, acyclic (no recurrent loops) graph whose nodes are the strongly connected components of *G*. Any neurons that were not connected to the dominant network were left out of our calculations.

Orderability was calculated as the fraction of neurons that were not included within any recurrent loop in *G* (i.e. the number of neurons that were their own strongly connected component of size 1, divided by the total number of neurons). Feedforwardness was calculated by examining each path in *G*_*C*_ that emanated from a “starting node” (a node with no incoming connections) and finished at an “ending node” (a node with no outgoing connections), dividing the number of nodes in that path by the total number of neurons represented by those nodes, then averaging this ratio over all paths from starting to ending nodes. Treeness was calculated by (1) calculating the forward path entropy *h*_*f*_(*v*_*i*_) of each starting node *v*_*i*_ in *G*_*C*_, (2) computing H_f_, the mean forward path entropy over all starting nodes, (3) calculating the backward path entropy *h*_*b*_(*u*_*i*_) of each ending node *u*_*i*_ in *G*_*C*_, (4) computing H_b_, the mean backward path entropy over all ending nodes, and (5) taking the normalized difference between the mean forward and backward path entropies, with the final feedforwardness given by (H_f_ - H_b_)/max(H_f_, H_b_). The forward path entropy of a starting node is the entropy of all paths emanating from that node and finishing at any ending node, *h*(*v*_*i*_) = -Σ_path_P(path) log_2_ P(path)) where P(path) is the probability of taking a given path by starting at *v*_*i*_ and repeatedly stepping to a downstream node (selecting uniformly at random among all possible downstream nodes) until an ending node is reached. The backward path entropy of an ending node, *h*(*u*_*i*_), is the equivalent quantity, but starting from the ending node and traversing the network backwards until a starting node is reached. See (Corominas-Murtra et al., 2013) for further mathematical details.

Fully random networks were generated using the same number of neurons as the empirical network and assigning connections at random with the same connection probability as measured empirically. Degree-constrained random networks were generated by fixing the number of incoming and outgoing connections of each neuron in the random network to match those of a corresponding neuron in the empirical network (with 1-to-1 correspondence), but otherwise assigning connections between neuron pairs at random.

### Neurotransmitter predictions

We predicted the neurotransmitter for 496 neurons within 35 cell types using the method presented in (Eckstein et al. 2020). We used FlyWire (Dorkenwald et al. 2020) to retrieve all automatically detected pre-synaptic sites in each neuron segment and filtered out synapses with a cleft score below 50 to remove likely false positives (Buhmann et al 2020). We predicted the neurotransmitter of each pre-synaptic site individually and defined the neurotransmitter of an entire neuron as the predicted majority neurotransmitter over its synapses. We labeled a neuron as inconclusive if its majority transmitter was predicted for less than 65% of its synapses. Using this cutoff leads to a greater than 95% prediction accuracy for the three fast-acting neurotransmitters GABA, acetylcholine and glutamate on the test set presented in (Eckstein et al. 2020) (see Supp. Fig. 9A). We validated whether neurotransmitter predictions from automatically detected pre-synaptic sites were robust by predicting the neurotransmitter of neurons within 6 cell types with known neurotransmitters and find perfect agreement for all neurons with conclusive neurotransmitter predictions. Neurotransmitter predictions for all cell types are shown in Supp. Fig. 9.

The classifier predictions for all but 5 cell types (pMN2/vpoDN, SAD_pr02, AVLP_pr04, A1, and SAD_pr01) agreed for at least 60% of individual neurons in each class. Therefore, we did not assign a neurotransmitter identity to these cell classes. The one exception was AVLP_pr04, in which we found evidence for multiple cell types. MCFO results revealed that the split-GAL4 line labeling AVLP_pr04 consisted of two cells per hemisphere, both with diffuse and widespread branching patterns across the AVLP and PVLP, but with one cell type having a projection toward the midline (Supp. Fig. 2). We identified EM reconstructions matching both these cell types and included them as AVLP_pr04 (Fig. 2). The cells with the midline projection in both hemispheres were predicted to be dopaminergic (Supp. Fig. 9C), whereas the two remaining cells were predicted to be GABAergic or glutamatergic. We also found that synaptic connectivity was different for the dopaminergic vs. non-dopaminergic cells (Fig. 7D). Therefore, we conclude that AVLP_pr04 contains one dopaminergic cell per hemisphere and one additional cell with inclusive neurotransmitter results.

### Calcium imaging

For calcium imaging experiments, we used 3-11 day-old virgin female flies reared at 25C and housed in groups of 1-8 flies/vial after eclosion. Calcium imaging experiments were performed during the fly’s light cycle. Flies were mounted and dissected as described in (Murthy and Turner, 2013) with the following modifications. Before mounting the fly, we removed the wings and legs with forceps. We removed fat covering the brain with forceps or with mouth suction through a sharp glass pipette. We ensured the aristae were intact and free by gently blowing on the fly before and after each experiment. We also looked for abdominal contractions in response to gentle blowing to indicate the fly was alive and healthy before and after each experiment. We monitored temperature and humidity with a thermometer and hygroscope (Traceable 15-077-963, Webster TX) placed on the air table with the two-photon microscope. Temperature and humidity were stable within an experiment, with fluctuations of <1C and <5% humidity across 8 hours of imaging (∼1 fly/hr).

Imaging was performed as previously described (Deutsch et al., 2019). Briefly, we used a custom-built two-photon laser scanning microscope controlled in MATLAB by ScanImage 5.1 (Vidrio). We imaged single planes at 8.5 Hz (256 x 256 pixels). Pixel size was 0.75 μm x 0.75 μm. After dissection, a fly was placed under the microscope with continuous saline perfusion delivered to the meniscus. Sound was delivered through an earbud speaker (Koss, 16 Ohm impedance; sensitivity, 112 dB SPL/1 mW), which was attached to a long thin tube (12cm, diameter: 1mm) placed ∼2mm from the fly’s head (directed toward the aristae) and controlled by custom software in MATLAB (Clemens et al., 2018b). Sound intensity was calibrated by measuring the sound particle velocity component for a range of frequencies (100-800 Hz). The detailed procedures and cross-calibration between the pressure and the pressure gradient microphone were described in (Göpfert et al., 2006). To estimate the sound amplitude of each stimulus we placed the calibrated gradient microphone at the same position as the fly (2-3 mm from sound tube outlet) in separate experiments. The recorded voltage was then converted to particle velocity (with units mm/s). The output signal was corrected according to the measured intensities to ensure equivalent intensity across frequencies, as previously described (Tootoonian et al., 2012).

### ROI selection

To decide which region of interest (ROI) to record from in each neuron, we first sampled from multiple ROIs, depending on the morphology of the neuron and on the level of baseline GCaMP6s expression (ie, only ROIs with some baseline level of GCaMP expression were visible under the two-photon microscope). In some lines, multiple ROIs were visible and responded relatively robustly. In those cases, we narrowed down which ROI to focus on based on response strength and ability to locate roughly the same ROI across flies. In other lines, baseline GCaMP expression was low, so we imaged from the ROI that was visible within a moderate laser power, with the goal of choosing roughly a similar ROI in each fly. We sampled from both the left and the right hemisphere within each line but from different flies. In all of the lines, responses occurred in both hemispheres, suggesting either that we stimulated both aristae or that ROIs at the level of the WED/VLP respond bilaterally.

### Stimulus generation and delivery

Sound stimuli were generated at a sampling frequency of 10kHz using custom MATLAB software and previously published techniques (Deutsch et al., 2019). To characterize a given neuron type as auditory or non-auditory, we used a stimulus set consisting of pulse song, sine song, and white noise, each with 10 sec duration and 10 sec pre- and post-stimulus silent periods. The intensity of pulse and sine stimuli was 5 mm/sec and the intensity of the white noise stimulus was 2 mm/sec. To further characterize auditory tuning, we used a stimulus set consisting of 5 pure tones at a range of frequencies (100, 200, 300, 500, and 800 Hz) and 5 pulse stimuli with a range of interpulse intervals (16, 36, 56, 79, and 96 ms), all at an intensity of 4 mm/sec. Each of these stimuli were 4 sec in duration, with 5 sec pre-stimulus and 10 sec post-stimulus silent periods. Within a stimulus set, a block was defined as one presentation of each stimulus. Stimulus order was randomized within each block, and 3-7 blocks were presented to each fly.

### Analysis

Data were analyzed using custom software in MATLAB. ROIs were drawn manually based on a z-projection of the td-Tomato signal. We calculated the mean fluorescence within each ROI frame-by-frame. If any stimulus block contained drastic, brief (1-3 sec) increases in fluorescence, indicative of wax or other particles moving into the imaged region, we discarded the entire block. We used only flies in which 3 or more blocks were available for analysis. We corrected for gradual, modest changes in fluorescence during each recording by de-trending the data. To detrend, we concatenated fluorescence traces over an entire recording, used a running percentile filter (20^th^ percentile, 50 sec window, 5 sec shift), and subtracted this long-term trend from the recording. For each stimulus, we calculated dF/F as (F(t)-F_0_)/F_0,_ where F_0_ was defined as the mean fluorescence during the 10 sec prestimulus silent period.

To quantify across-fly variability within a spit-GAL4 line, we measured Euclidean distances between z-scored responses to pulse, sine, and white noise between every possible pair of flies imaged from a given line. For each fly, we calculated the mean calcium trace elicited by pulse, sine and, white noise stimuli, and then concatenated the z-scored mean traces in that order before measuring Euclidean distances.

### Stimulus modulation of calcium signals

To determine whether the fluorescence within a given ROI was modulated by at least one of the acoustic stimuli, we followed the method previously described in (Pacheco et al., 2020). Briefly, we modeled the GCaMP fluorescence trace as a convolution of the stimulus history and a set of filters, one per stimulus (ie, 3 filters for the screen stimuli and 10 filters for the frequency/IPI tuning stimuli), with the filter duration matching the duration of the stimulus. To estimate each filter, we used 80% of the stimulus repetitions as training data and the remaining 20% of the repetitions as test data. Filters were estimated using ridge regression (Park and Pillow, 2011). We convolved the estimated filters with the stimulus history to generate the predicted signal for each ROI. All possible combinations of training and test data repetitions were used, which gave a total of 3-15 predicted signals (3-6 total repetitions, respectively) per ROI. We then measured the Pearson correlation coefficient between the raw (ie, test repetitions) and the predicted signals. We used bootstrapping to determine the statistical significance of the resulting correlation coefficients. We randomly shuffled each test fluorescence trace in 10 sec bins, and then calculated the Pearson correlation coefficient between each of 10,000 independently shuffled test signals and the predicted signal. P-values for the correlation coefficients were defined as the fraction of shuffled correlation coefficients > 30^th^ percentile of the estimated correlation coefficients. P-values were corrected for multiple comparisons with the Benjamini-Hochberg procedure, with a false detection rate of 0.05.

### Analysis of auditory responses

We measured the strength of calcium responses to auditory stimuli only in ROIs which we determined to be statistically modulated by the stimuli. In these cases, we calculated the integral of the dF/F signal starting with stimulus onset and ending 4 sec after stimulus offset, and then divided by the total time of the integral window.

We used the integrals elicited by pulse and sine during the screen stimuli to assess each fly’s song mode preference. We calculated a song mode preference index as (pulse - sine) / (pulse + sine) after setting all negative integrals to 0. If both integrals were negative, reflective of net inhibition to both stimuli, we set the song mode preference to 0 to indicate that the response did not prefer either stimulus. Next we reasoned that in a pulse-preferring neuron, the pulse integral should be at least 250% of the sine integral, which corresponds to a preference index of 0.43 (Fig. 3D). Likewise, we required a sine-preferring neuron to have a sine response that was at least 250% of the pulse integral, which corresponds to a preference index of -0.43. To classify the preference of each cell type, we used the mean preference index across flies. Neurons with a mean preference index between -0.43 and 0.43 were classified as having intermediate preference.

### Hierarchical clustering of auditory responses

We clustered responses to pulse, sine, and white noise stimuli according to previously published methods (Pacheco et al., 2020). Briefly, we first calculated the median dF/F trace from each fly across stimulus repetitions, and concatenated the traces starting with pulse and ending with white noise (including pre- and post-stimulus periods). We then z-scored this trace for each fly. Next we hierarchically clustered these traces based on Euclidean distances and inner square distance metric between clusters (Ward’s method). To determine the number of clusters, we set a lower limit of three responses per cluster, which resulted in 13 total clusters across all our responses.

### Tuning curve generation and classification

To generate frequency and interpulse interval tuning curves, we measured the integrals in response to each stimulus. Next we classified frequency and interpulse interval tuning curve types based on previously published methods (Baker and Carlson, 2014; Groh et al., 2003). Briefly, if the minimum tuning curve value was >85% of the maximum tuning curve value, we classified the tuning as all-pass (ie, untuned to the tested stimulus parameter). For all other tuning curves, we fit both a sigmoid and a Gaussian function. If the *r*^2^ of both the sigmoid and Gaussian fits were <0.5, we classified the tuning as complex. Since high- and low-pass tuning curves can be well fit by both a Gaussian and a sigmoidal curve, we used the *r*^2^_sigmoid_/*r*^2^_Gaussian_ ratio. If *r*^2^_sigmoid_/*r*^2^_Gaussian_ was <0.85, we classified the tuning curve as band-pass if the Gaussian amplitude was positive and band-stop if the Gaussian amplitude was negative. For frequency tuning curves, if *r*^2^_sigmoid_/*r*^2^_Gaussian_ was≥ 0.85, we classified the tuning as high-pass if the ratio of the sigmoid slope to the sigmoid amplitude was positive, and low-pass if this ratio was negative. For IPI tuning curves, if *r*^2^_sigmoid_/*r*^2^_Gaussian_ was≥ 0.85, we classified the tuning as long-pass if the ratio of the sigmoid slope to the sigmoid amplitude was positive, and short-pass if this ratio was negative.

