## Supplementary material for "Neural Network Organization for Courtship Song Feature Detection in *Drosophila*": Tables 1 and 2

**Table 1. Neuropil abbreviations**

| Abbreviation | Neuropil |
|---|---|
| AL | antennal lobe |
| AME | accessory medulla |
| AMMC | antennal mechanosensory and motor center |
| AOTU | anterior optic tubercle |
| ATL | antler |
| AVLP | anterior ventrolateral protocerebrum |
| BU | bulb |
| CAN | cantle |
| CRE | crepine |
| EB | ellipsoid body |
| EPA | epaulette |
| FB | fan-shaped body |
| FLA | flange |
| GA | gall |
| GNG | gnathal ganglion |
| GOR | gorget |
| IB | inferior bridge |
| ICL | inferior clamp |
| IPS | inferior posterior slope |
| LAL | lateral accessory lobe |
| LH | lateral horn |
| LO | lobula |
| LOP | lobula plate |
| MB-CA | mushroom body calyx |
| MB-ML | mushroom body medial lobe |

| | |
|---|---|
| MB-PED | mushroom body pedunculus |
| MB-VL | mushroom body vertical lobe |
| ME | medulla |
| NO | nodulus |
| PB | protocerebral bridge |
| PLP | posterior lateral protocerebrum |
| PRW | prow |
| PVLP | posterior ventrolateral protocerebrum |
| SAD | saddle |
| SCL | superior clamp |
| SIP | superior intermediate protocerebrum |
| SLP | superior lateral protocerebrum |
| SMP | superior medial protocerebrum |
| SPS | superior posterior slope |
| VES | vest |
| WED | wedge |

**Table 2. Split-GAL4 IDs and Cell Type names**

| Split-GAL4 line ID | Existing name from literature or WED/VLP cell class name based on primary neuropil targeted |
|---|---|
| SS16374 | AVLP_pr01 |
| SS17963 | AVLP_pr03 |
| SS21914 | AVLP_pr04 |
| SS23281 | AVLP_pr05 / AVLP-PN1 (Dolan et al., 2019) |
| SS23627 | AVLP_pr06 |
| SS27880 | AVLP_pr07 |
| SS27883 | AVLP_pr08 |
| SS27885 | AVLP_pr02 |
| SS27919 | WED-VLP (Dorkenwald et al., 2020) / IVLP-VLP (Lai et al., 2012) |
| SS27932 | IPS_pr01/ WED-PN1 (Dolan et al., 2019) |
| SS27936 | WED_pr01 |
| SS27953 | AVLP_pr09 |
| SS28822 | AVLP_pr10 |
| SS29136 | AVLP_pr11 |
| SS29146 | AVLP_pr12 |
| SS35318 | AVLP_pr13 |
| SS35456 | AVLP_pr14 |
| SS35471 | AVLP_pr15 |
| SS35473 | A2 (Lai et al., 2012) |
| SS35484 | AVLP_pr16 |
| SS35488 | SLP_pr01 |
| SS35489 | AVLP_pr17 |
| SS35502 | AVLP_pr18 |
| SS35504 | PLP_pr01 |
| SS35916 | WED_pr02 |

| | |
|---|---|
| SS35920 | IC/SPS_pr01 |
| SS35944 | AVLP_pr19 |
| SS35996 | AVLP/PVLP_pr01 |
| SS36007 | GNG_pr01 |
| SS36012 | IPS_pr02 |
| SS36521 | PVLP_pr01 |
| SS36541 | AVLP_pr20 |
| SS36960 | AVLP_pr21 |
| SS36971 | SIP_pr01 |
| SS36978 | B1 (Kamikouchi et al., 2009; Lai et al., 2012) / aPN1 (Vaughan et al., 2014) |
| SS37007 | AVLP_pr22 |
| SS38918 | AVLP_pr23 / AV2 (Clemens et al., 2015) |
| SS38928 | PVLP_pr02 |
| SS38958 | IPS_pr03 |
| SS38981 | B2 (Kamikouchi et al., 2009; Tootoonian et al., 2012) |
| SS40303 | IPS/WED_pr01 |
| SS40316 | PVLP_pr03 |
| SS40354 | AVLP_pr24 |
| SS40361 | AVLP_pr25 |
| SS40366 | AVLP_pr26 |
| SS41342 | AVLP_pr27 |
| SS41376 | AVLP_pr28 |
| SS41378 | AVLP_pr29 |
| SS41685 | AVLP_pr30 |
| SS41704 | PLP_pr02 |
| SS41728 | AVLP_pr31 |
| SS41730 | AVLP_pr32 |
| SS41739 | SAD_pr01 |

| | |
|---|---|
| SS43299 | SAD_pr02 |
| SS43321 | SAD/AMMC_pr01 |
| SS43460 | AVLP_pr33 |
| SS43466 | SMP_pr01 |
| SS44510 | IB_pr01 |
| SS44818 | AVLP_pr34 |
| SS45858 | AVLP_pr35 |
| SS47261 | AVLP_pr36 |
| SS47509 | AVLP_pr37 |
| SS47517 | WED_pr03 |
| SS47537 | PVLP_pr04 |
| SS49115 | AVLP_pr38 |

**Table 2.**
