## Supplemental Figures and Legends for "Neural Network Organization for Courtship Song Feature Detection in *Drosophila*"

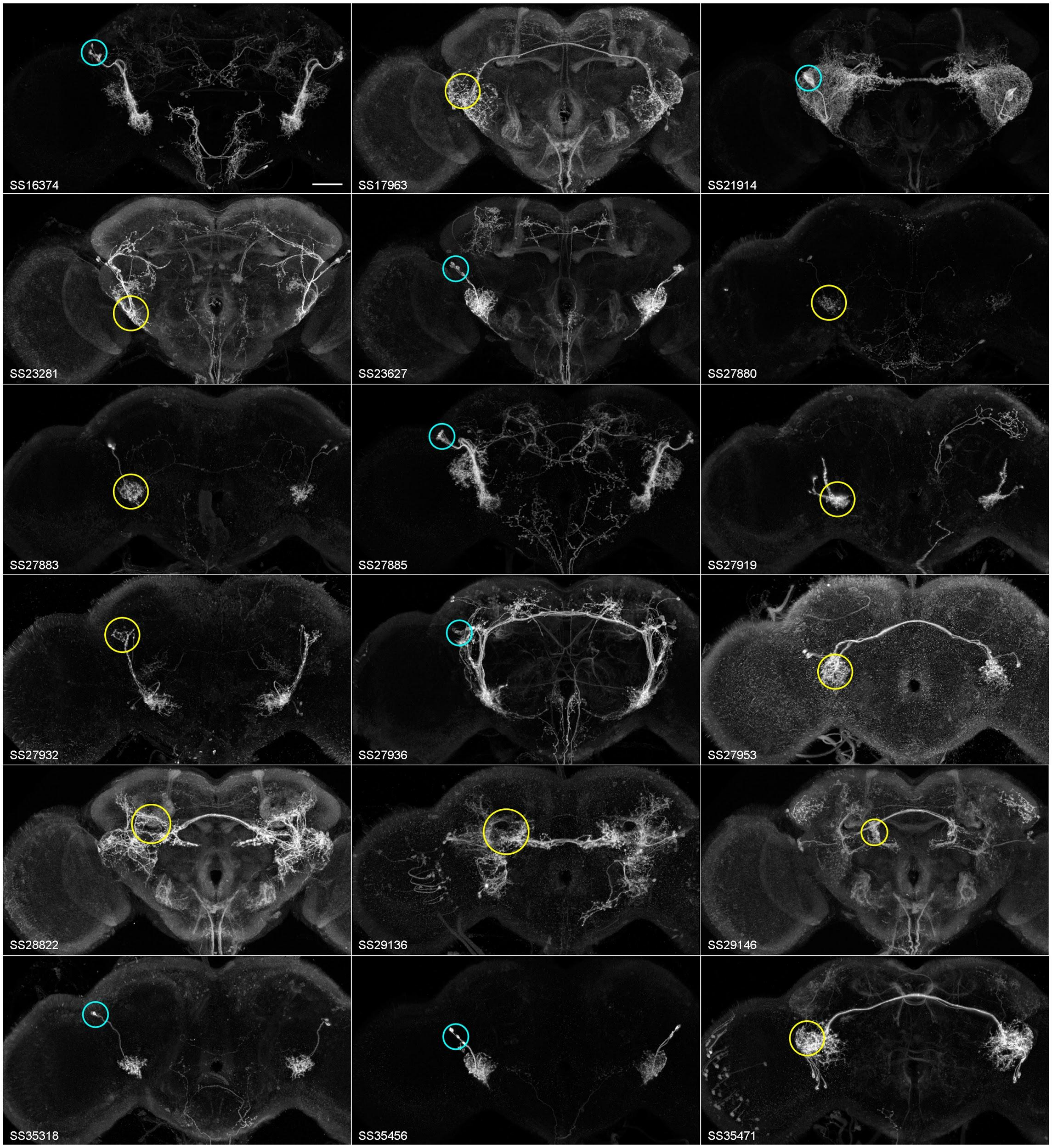


**Supplemental Figure 1. Expression patterns of the WED/VLP split-GAL4 collection.** Frontal maximum projections of split-GAL4 expression patterns in aligned brains for each line of the collection (split-GAL4 line number in bottom left of each panel). The region of interest (ROI) imaged in each line is circled (cyan for somas, yellow for projections) in one hemisphere, although we sampled from both hemispheres across flies. For genotypes, see Methods: Fly Stocks. Scale bar: 40 microns.

Supplemental Figure 1 continued

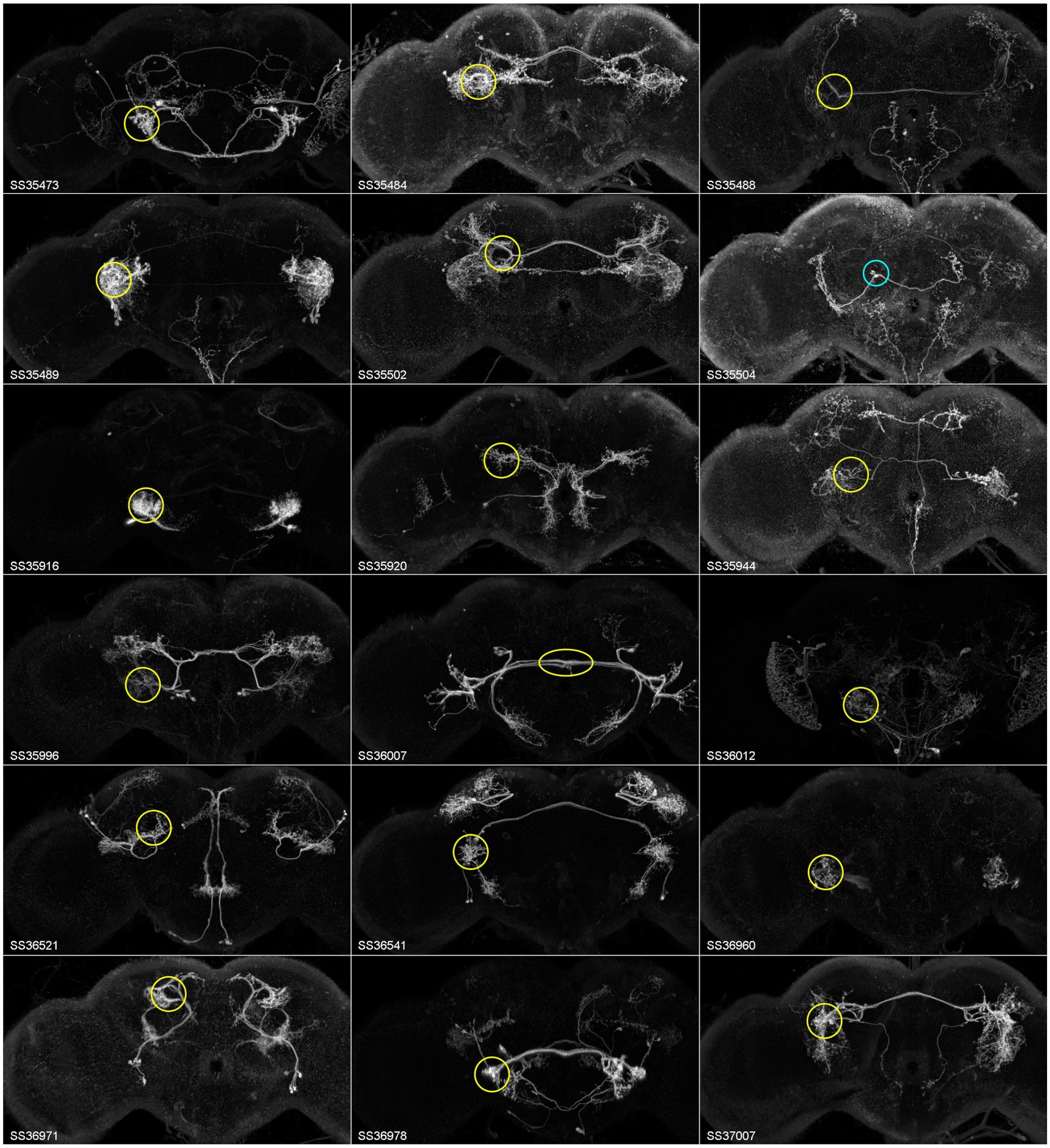

Supplemental Figure 1 continued

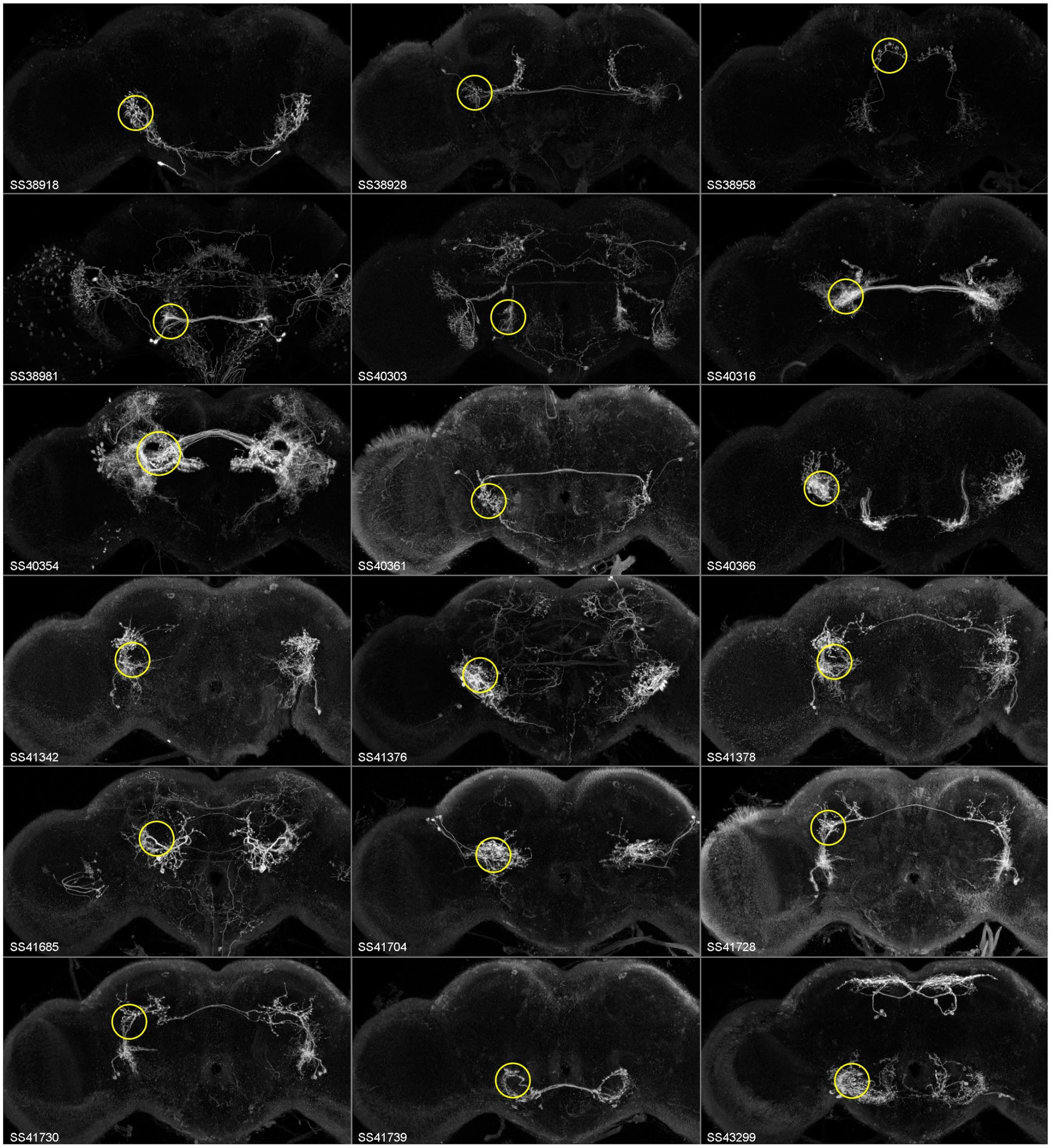

Supplemental Figure 1 continued

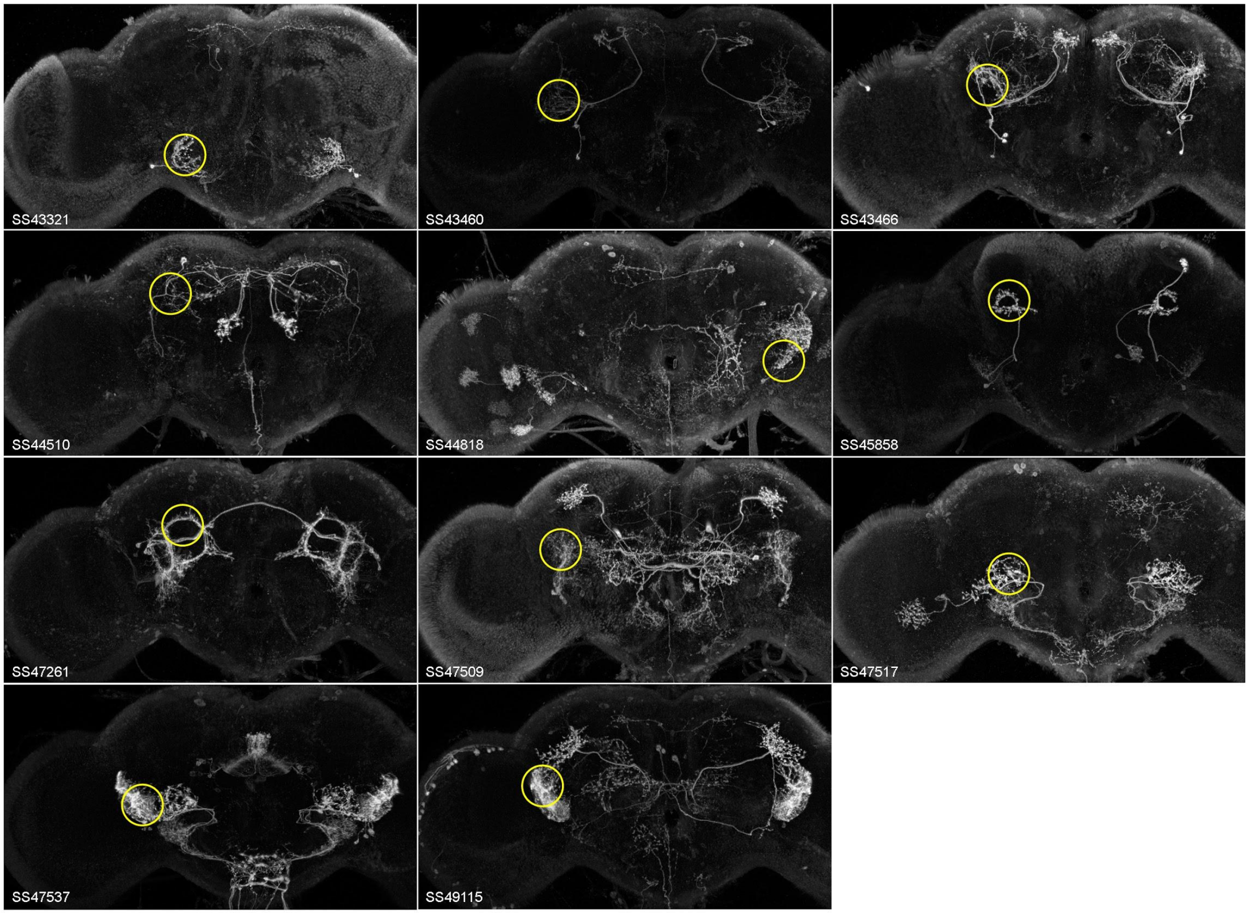

Supplemental Figure 2

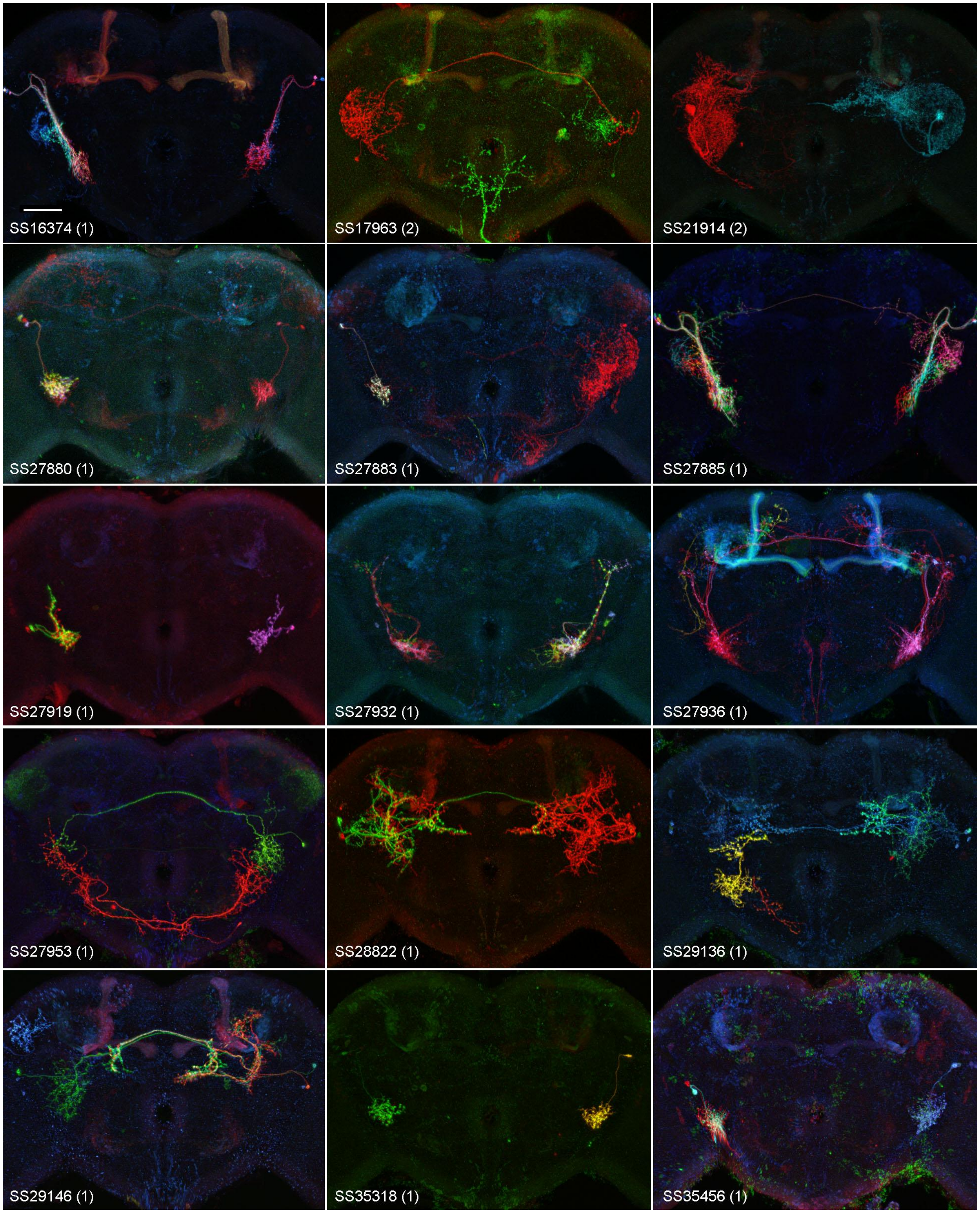


**Supplemental Figure 2. Stochastic labeling uncovers some cell type diversity within WED/VLP split-GAL4 lines.** Frontal maximum projections of aligned central brains with multicolor flip-out (MCFO) (Nern et al., 2015) expression labels individual cells in random colors and reveals different cell types that present in a split-GAL4 line. Composite images from 1-3 individuals (numbers in brackets on panels) to show in a compact fashion some of the cell types in each line. We erred on the side of including not just the typical cell types but also contaminating cell types, even if they were rare. Since only a few flies (approx. 8) of each line were stained with this stochastic method, a small number of lines showed no cells in the auditory areas (by chance) and were excluded from this figure. For a few cell classes, MCFO images revealed commissural and non-commissural neurons that were otherwise morphologically similar. For instance, the non-commissural neurons of SS27885 and SS41728 were very similar to those of SS16374 and SS41730, respectively. Therefore, we split these groups into different cell types (AVLP_pr01 and AVLP_pr32 are the non-commissural neurons, and AVLP_pr02 and AVLP_pr31 are the commissural neurons, respectively) (Fig. 2), and imaged from each cell type within different lines. SS47261 contained both commissural and non-commissural neurons, but since these neurons only appeared in one split-GAL4 line, we grouped the commissural and non-commissural neurons together as AVLP_pr36 (Fig. 2). Scale bar: 40 microns.

Supplemental Figure 2 continued

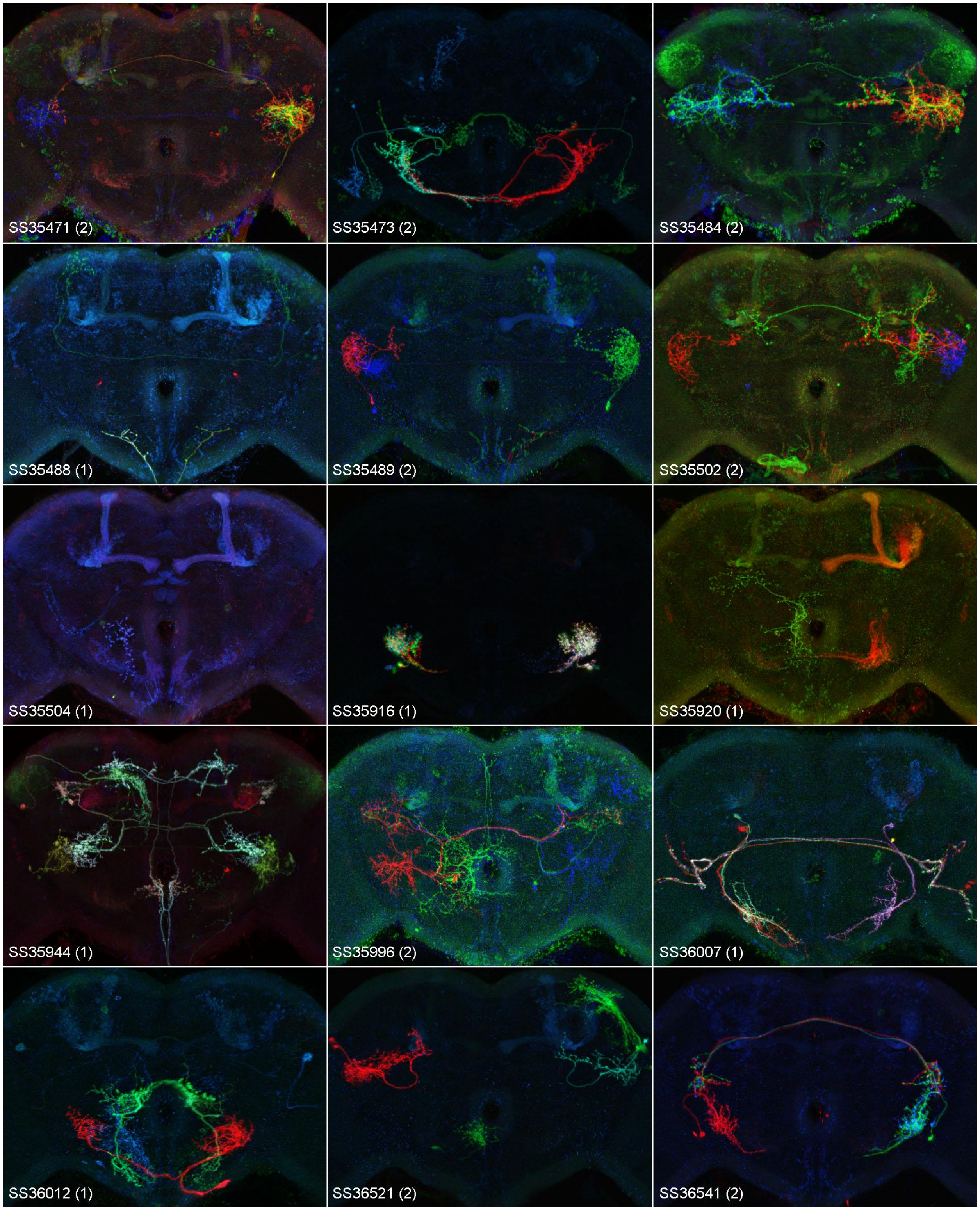


Supplemental Figure 2 continued

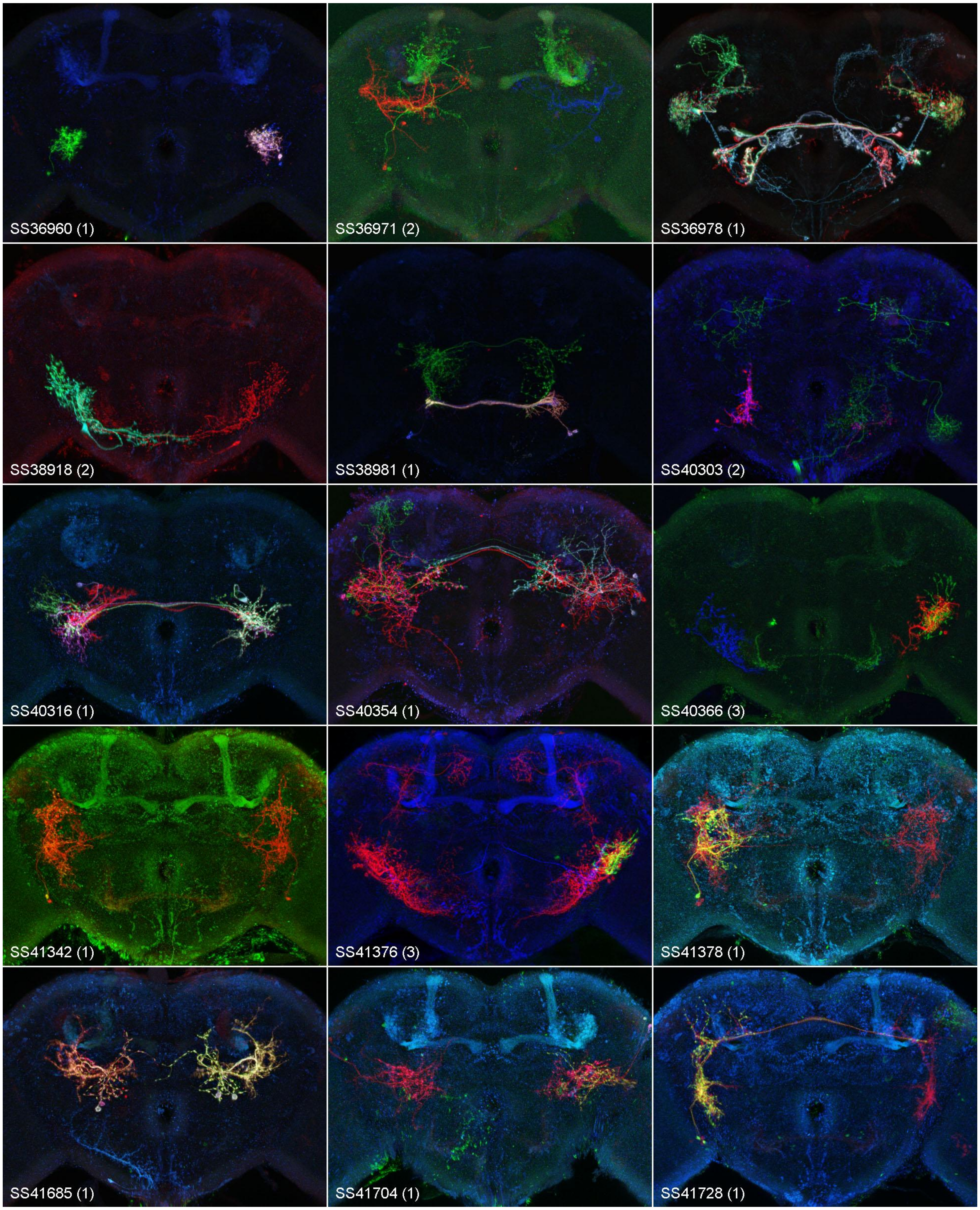


Supplemental Figure 2 continued

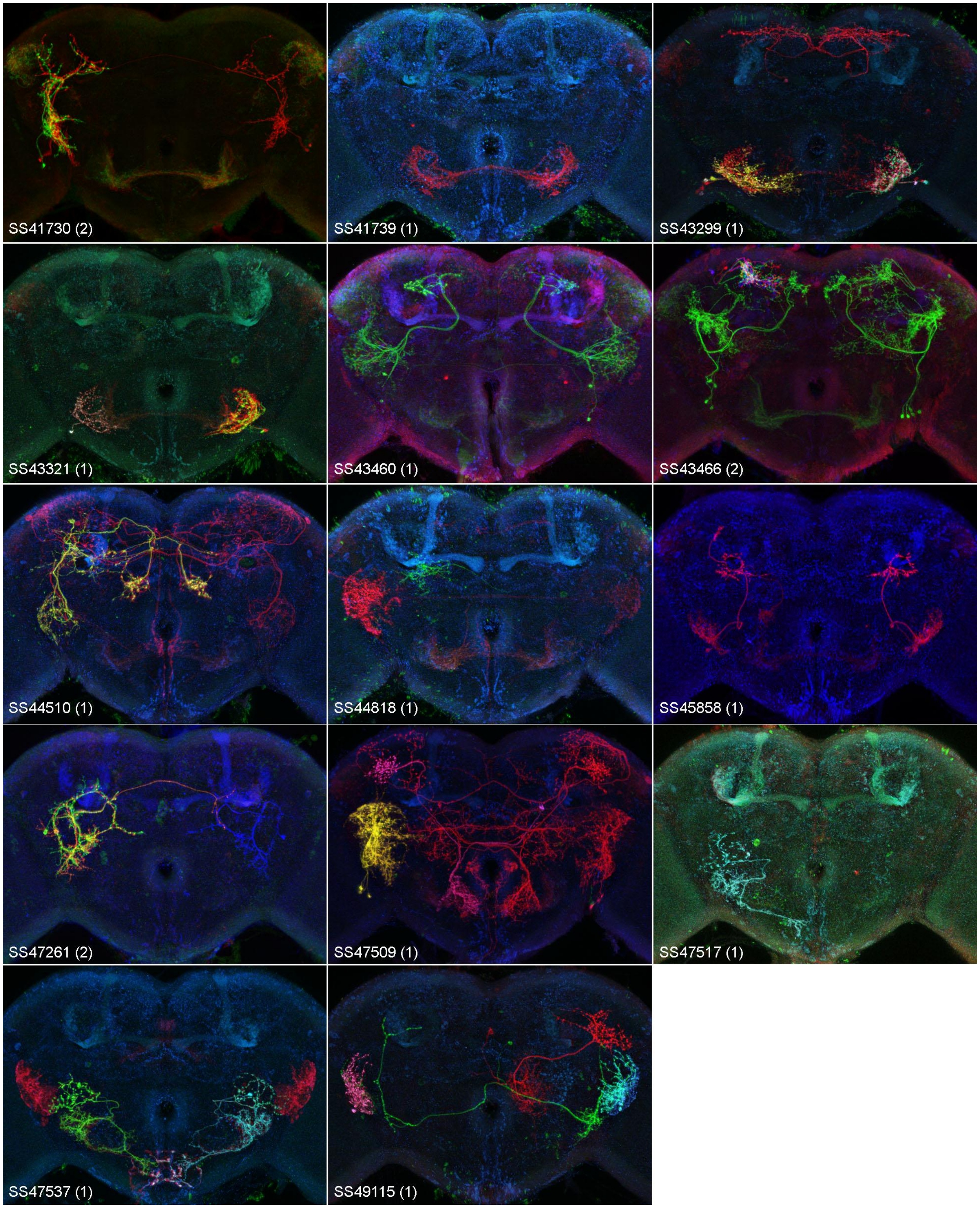
SS41730 (2)
SS41739 (1)
SS43299 (1)
SS43321 (1)
SS43460 (1)
SS43466 (2)
SS44510 (1)
SS44818 (1)
SS45858 (1)
SS47261 (2)
SS47509 (1)
SS47517 (1)
SS47537 (1)
SS49115 (1)

Supplemental Figure 3

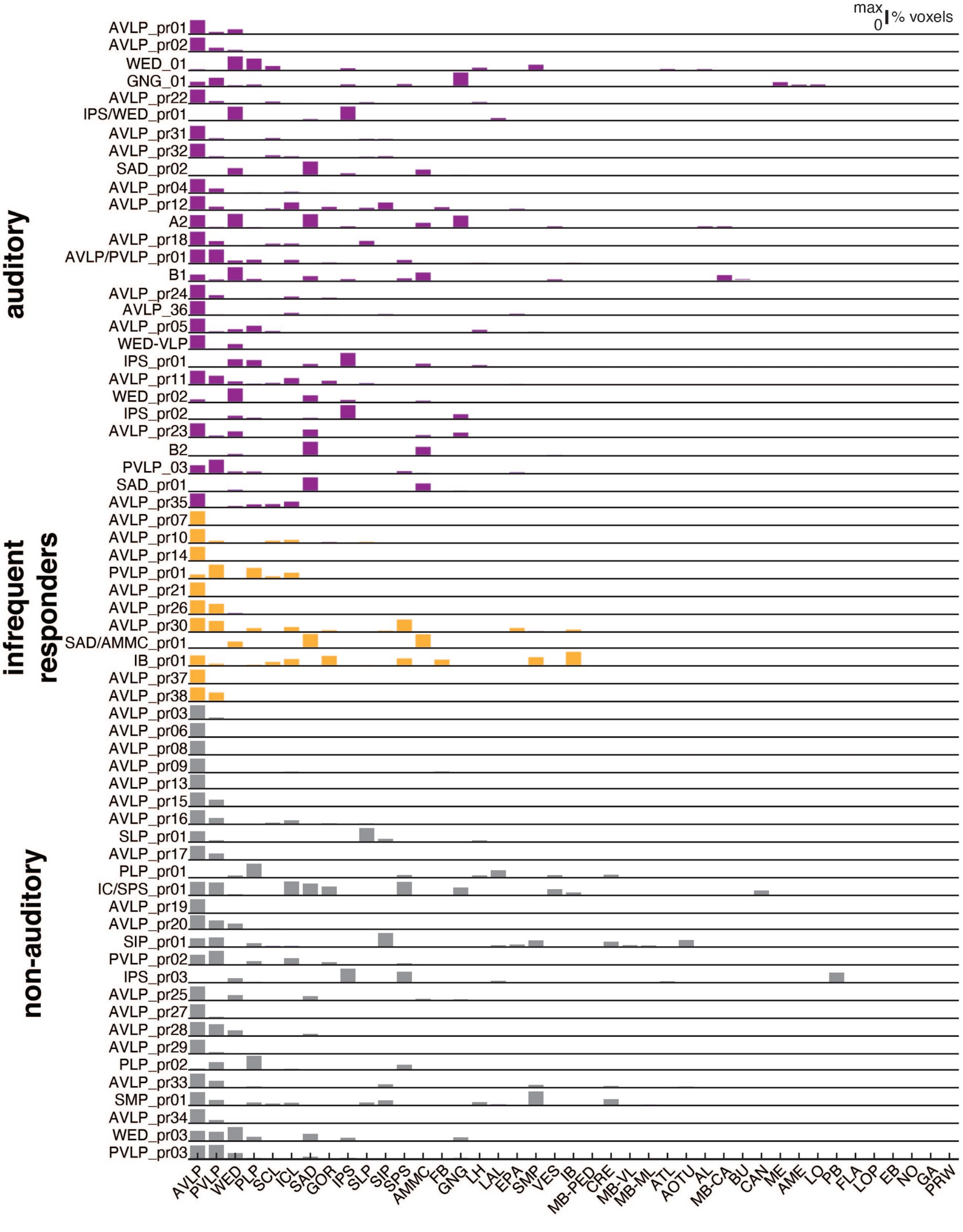

**Supplemental Figure 3. Neuropil innervation (by volume) for each WED/VLP cell type.** Histograms of the percentage of each cell class's processes in each brain neuropil (see Table 1 for neuropil names; see Methods). WED/VLP neurons within each split-GAL4 line were named based on the neuropil with the highest density of processes. If a cell class had equivalent processes in two neuropils, both neuropils were used in the name. Each cell class's histogram is normalized to its maximum.

Supplemental Figure 4

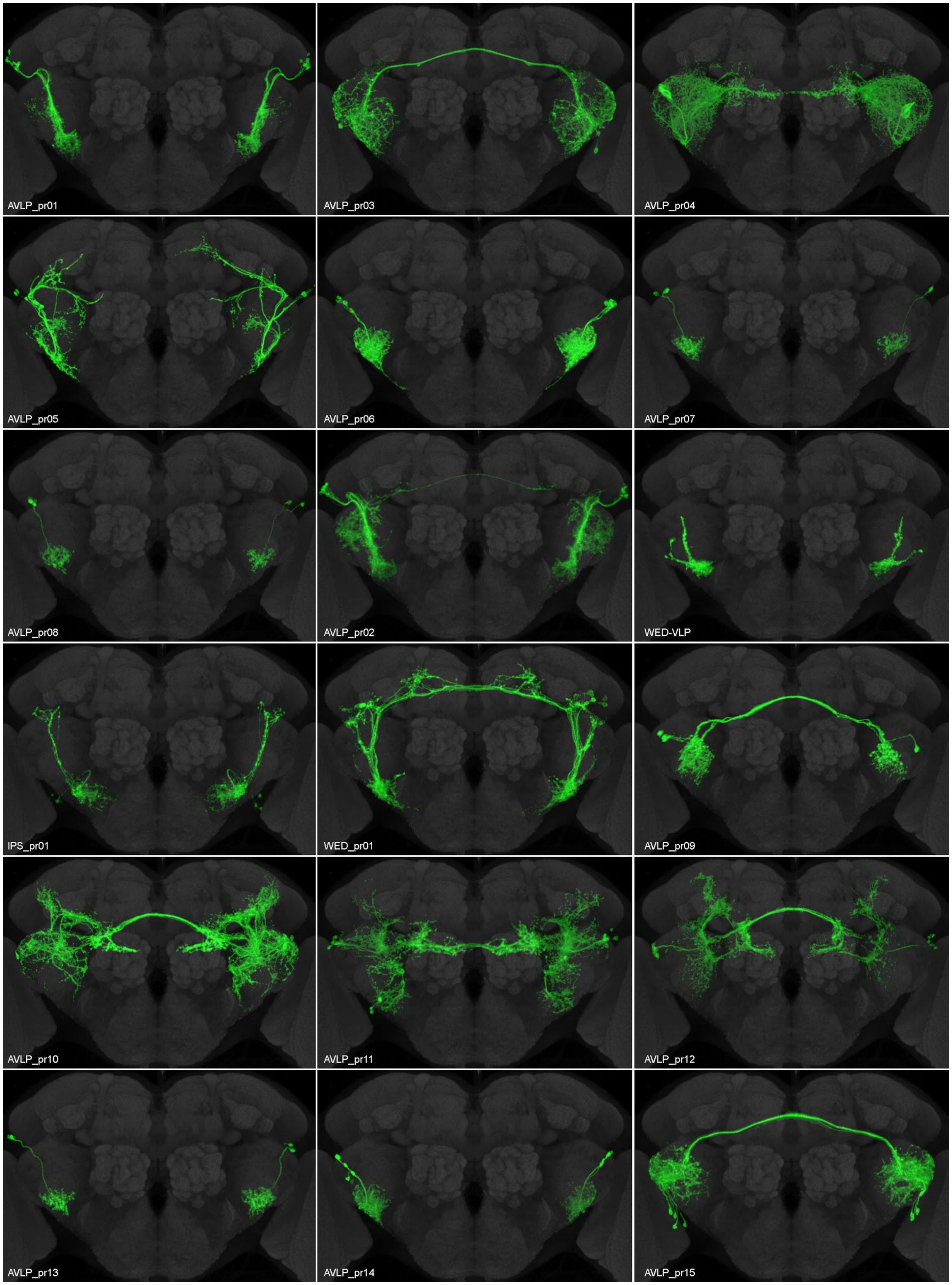

**Supplemental Figure 4. Segmented cell classes of the WED/VLP split-GAL4 collection.** Aligned central brains with split-GAL4 expression (from Supp. Fig. 1) patterns manually segmented to show the prominent cell class(es) (green) in the WED and/or VLP. The targeted cell class in each split-GAL4 line was named based on which neuropil had the highest density of projections (see Methods). Gray: nc82. In some brains (e.g., AVLP_pr11), cell types overlapped too much in 3D to be separated, however cell types could often be distinguished with the aid of stochastic labeling (Supp. Fig. 2) images.

### Supplemental Figure 4 continued

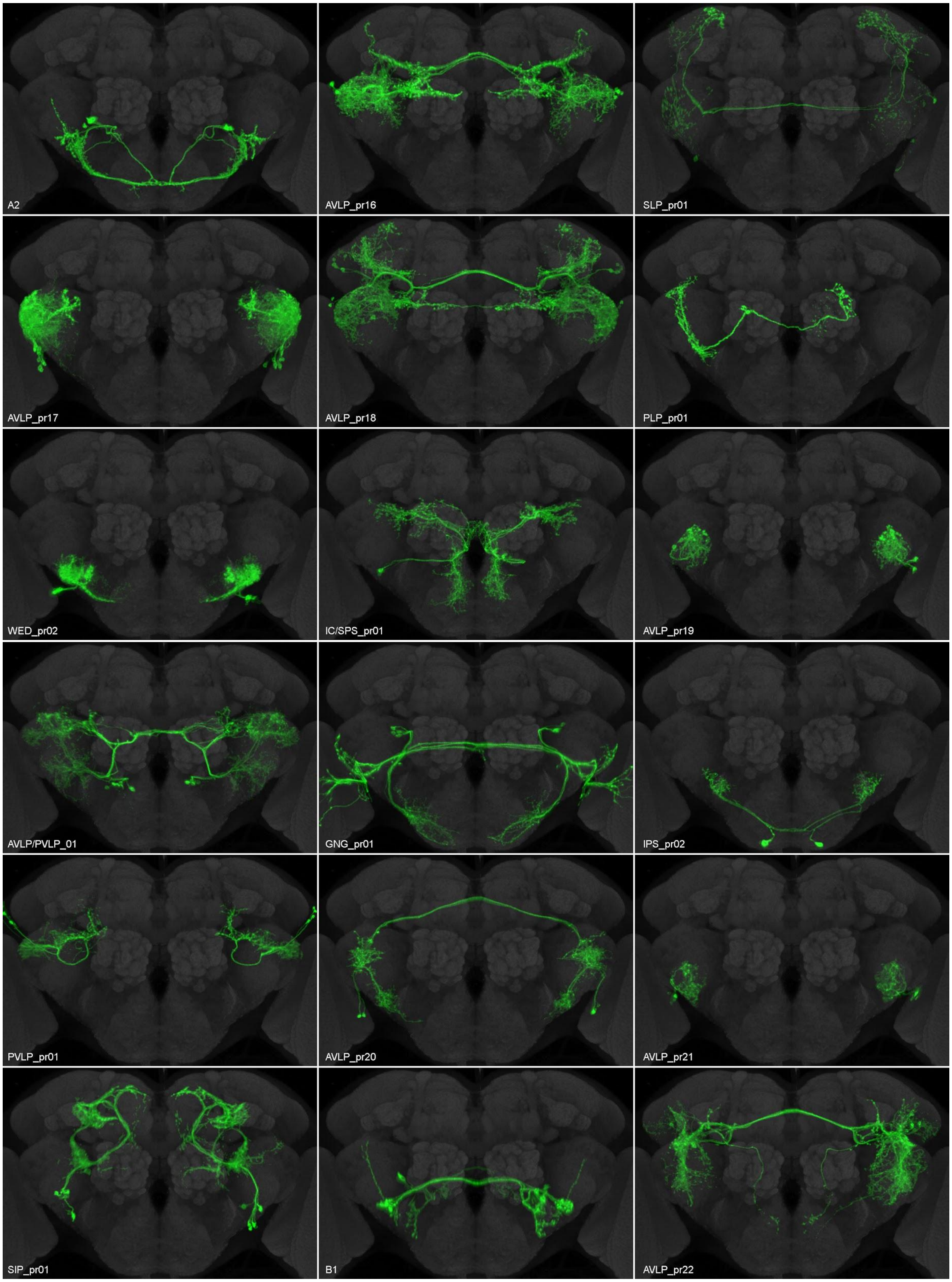

Supplemental Figure 4 continued

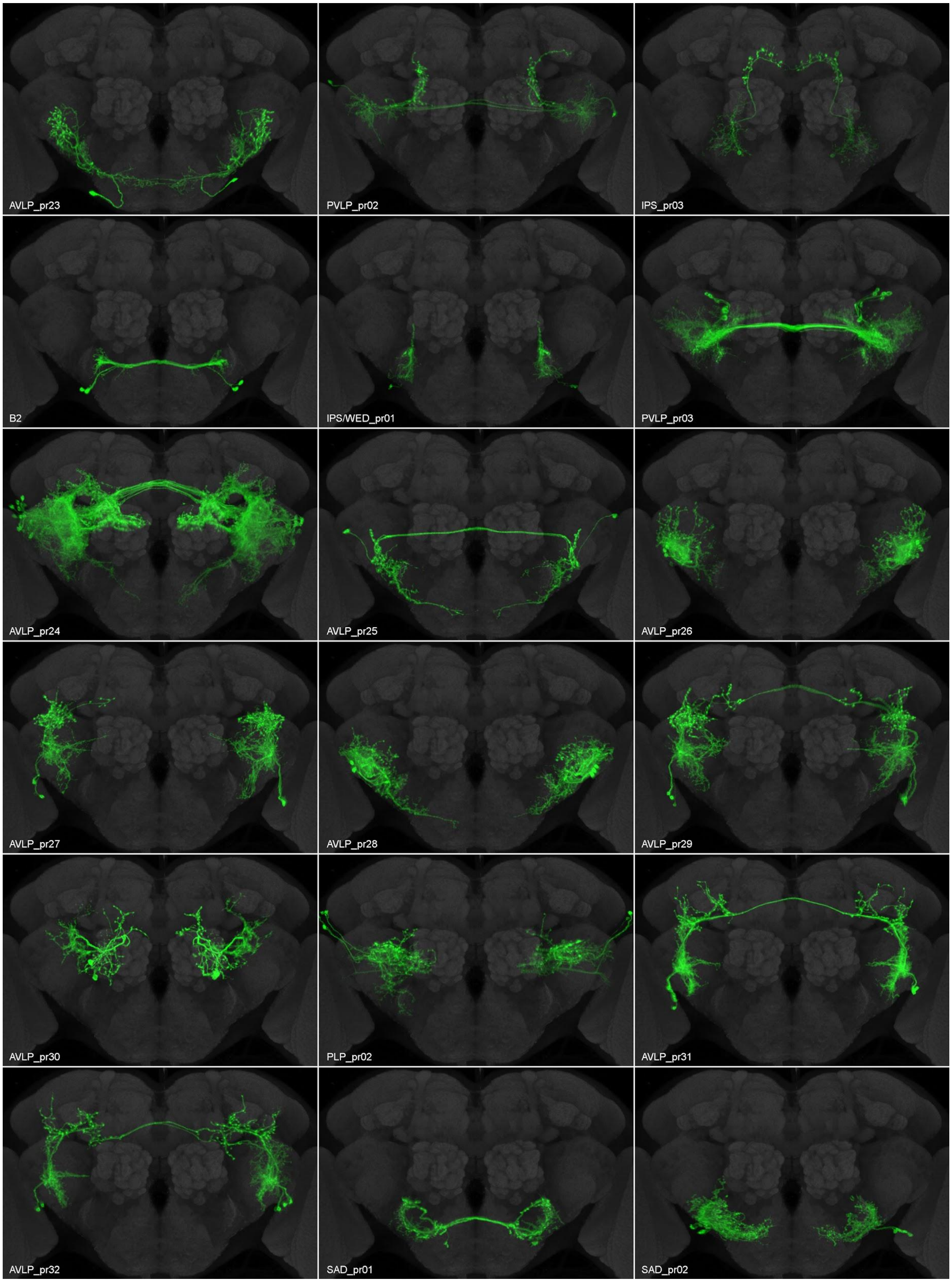

Supplemental Figure 4 continued

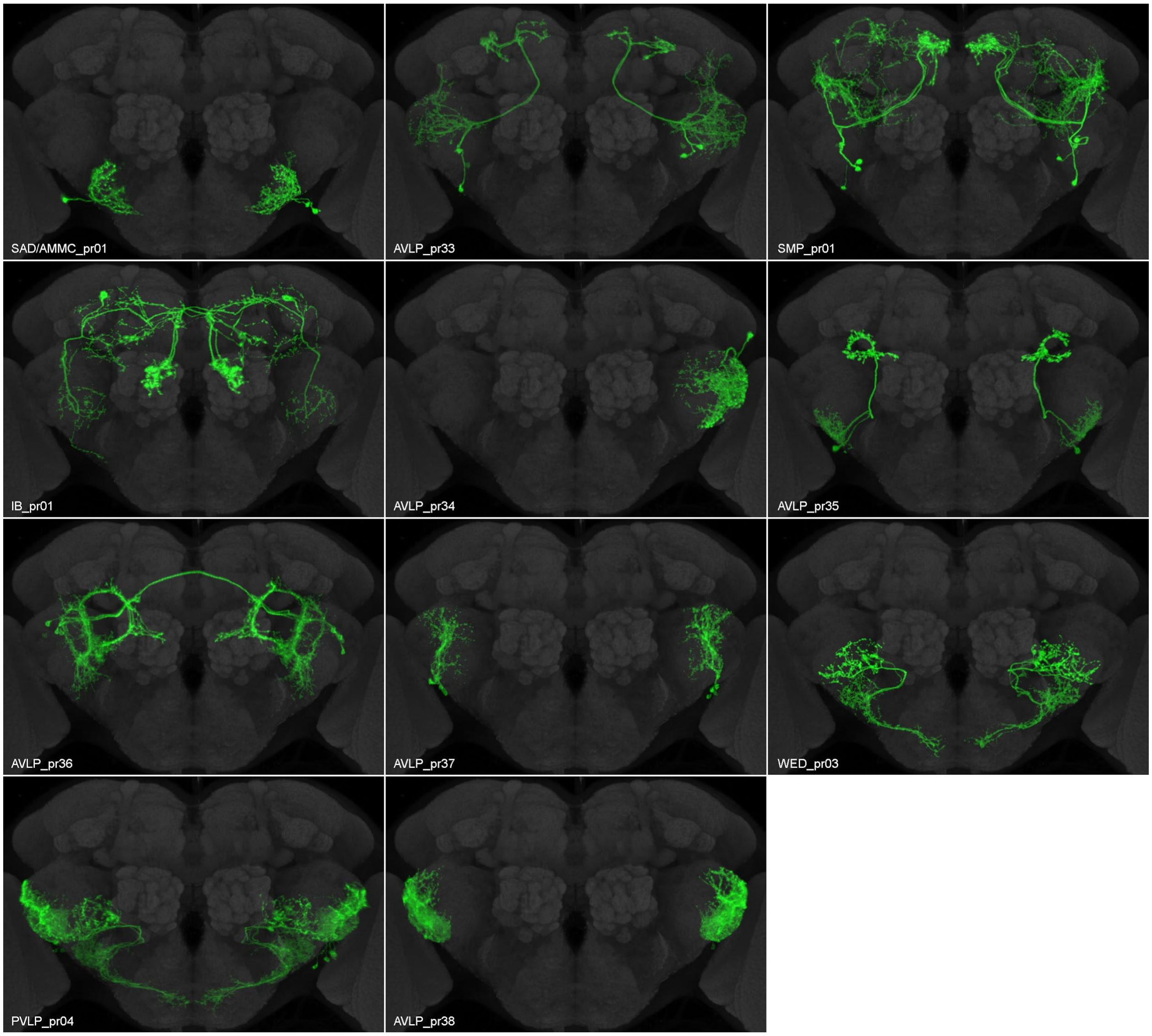

Supplemental Figure 5

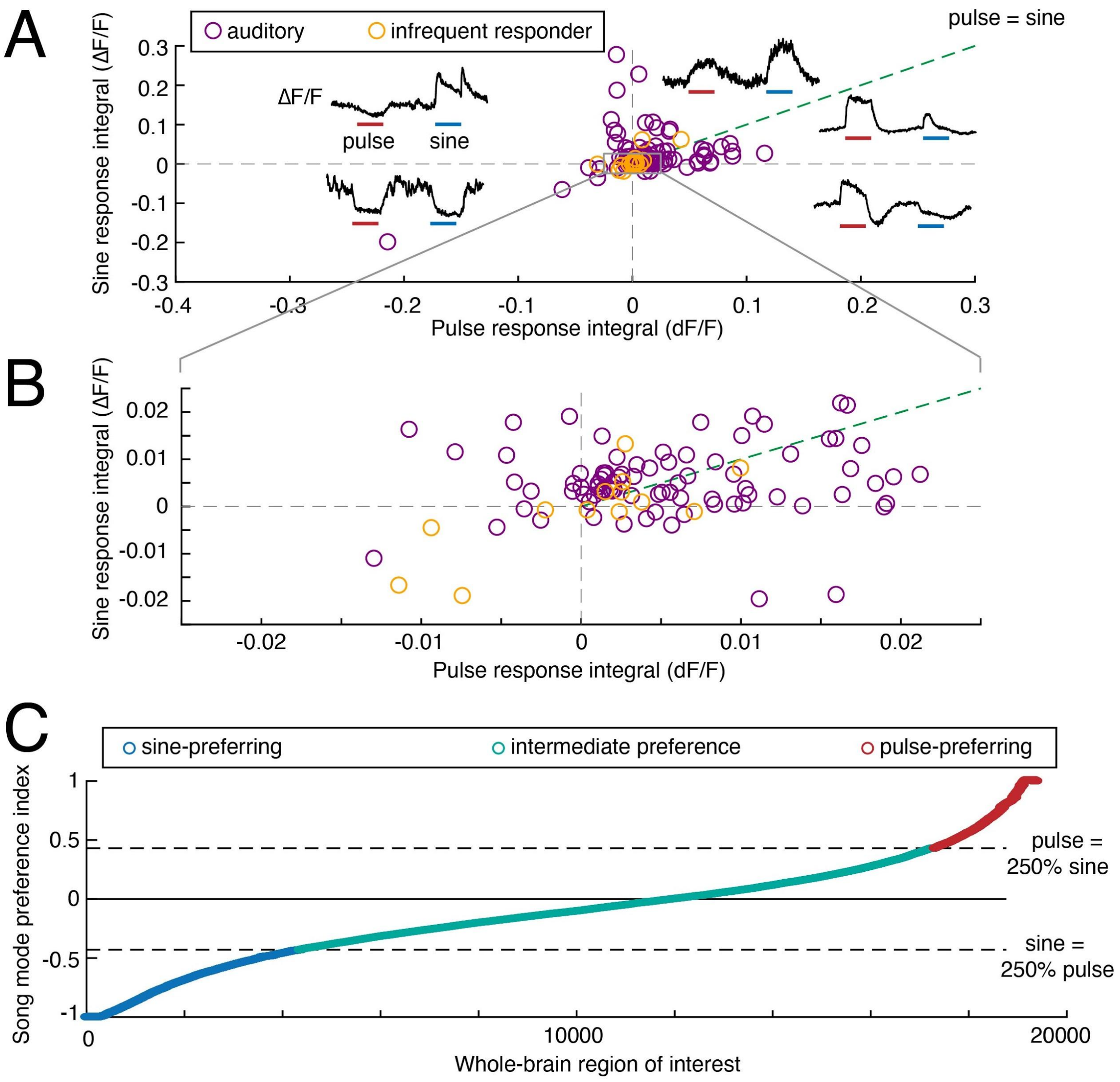


**Supplemental Figure 5. Measuring selectivity for pulse versus sine stimuli across WED/VLP neurons and the entire central brain.** A) The integral of each trial-averaged response (open circles) to pulse vs. the integral response to sine stimuli. Black box indicates region enlarged in (B). Each dot represents the responses from one fly. (C) The song mode preference index for 19,389 auditory-responsive regions of interest (ROIs) from the entire central brain, obtained via pan-neuronal imaging in a previous study (Pacheco et al., 2020).

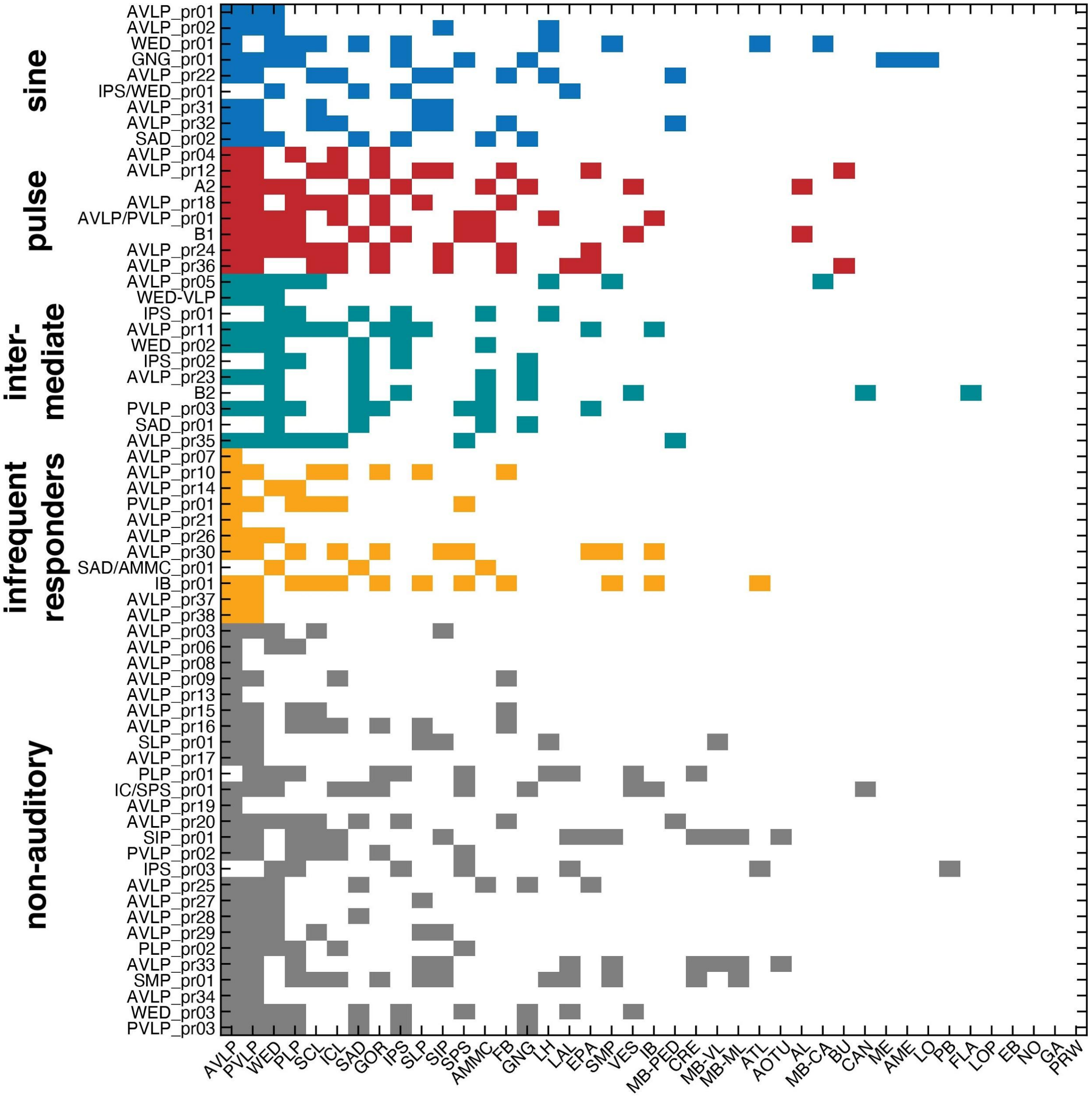


**Supplemental Figure 6. WED/VLP neurons send projections throughout the brain, independent of auditory responses.** A) Neuropils innervated by WED/VLP neuron classes, defined as having at least 1% of a cell class's total volume in each neuropil. Each WED/VLP neuron type is colored according to its overall sine or pulse preference (see Fig. 3).

Supplemental Figure 7

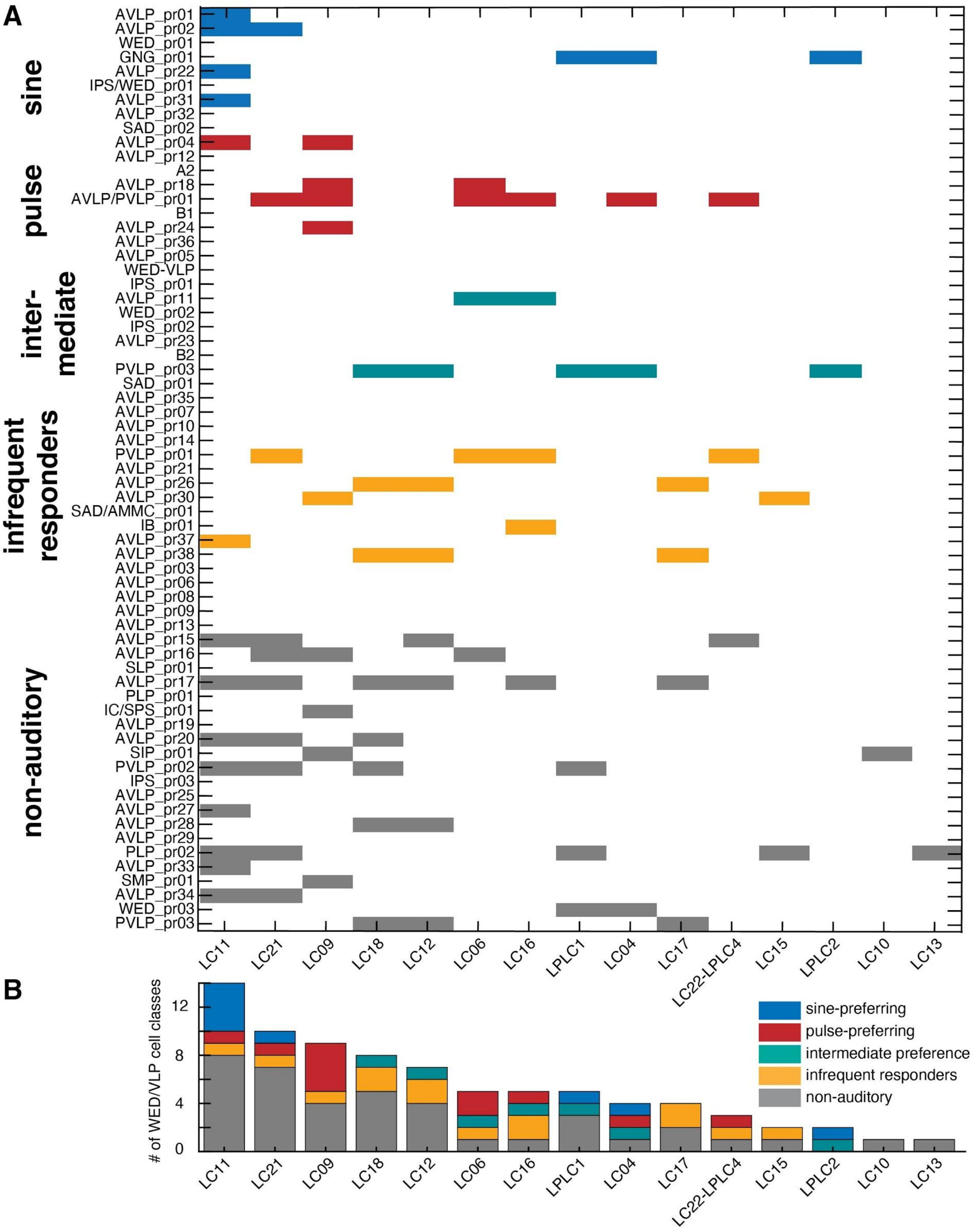

**Supplemental Figure 7. WED/VLP neurons with projections in optic glomeruli.** A) Optic glomeruli innervated by WED/VLP cell classes (see Methods), defined as having at least 1% of a cell class's total volume in each glomerulus. Only optic glomeruli that overlap with at least one WED/VLP cell class are shown. Each WED/VLP neuron type is colored according to its overall sine or pulse preference (see Fig. 3). B) Histogram of the total number of WED/VLP cell classes with innervation in each glomerulus.

Supplemental Figure 8

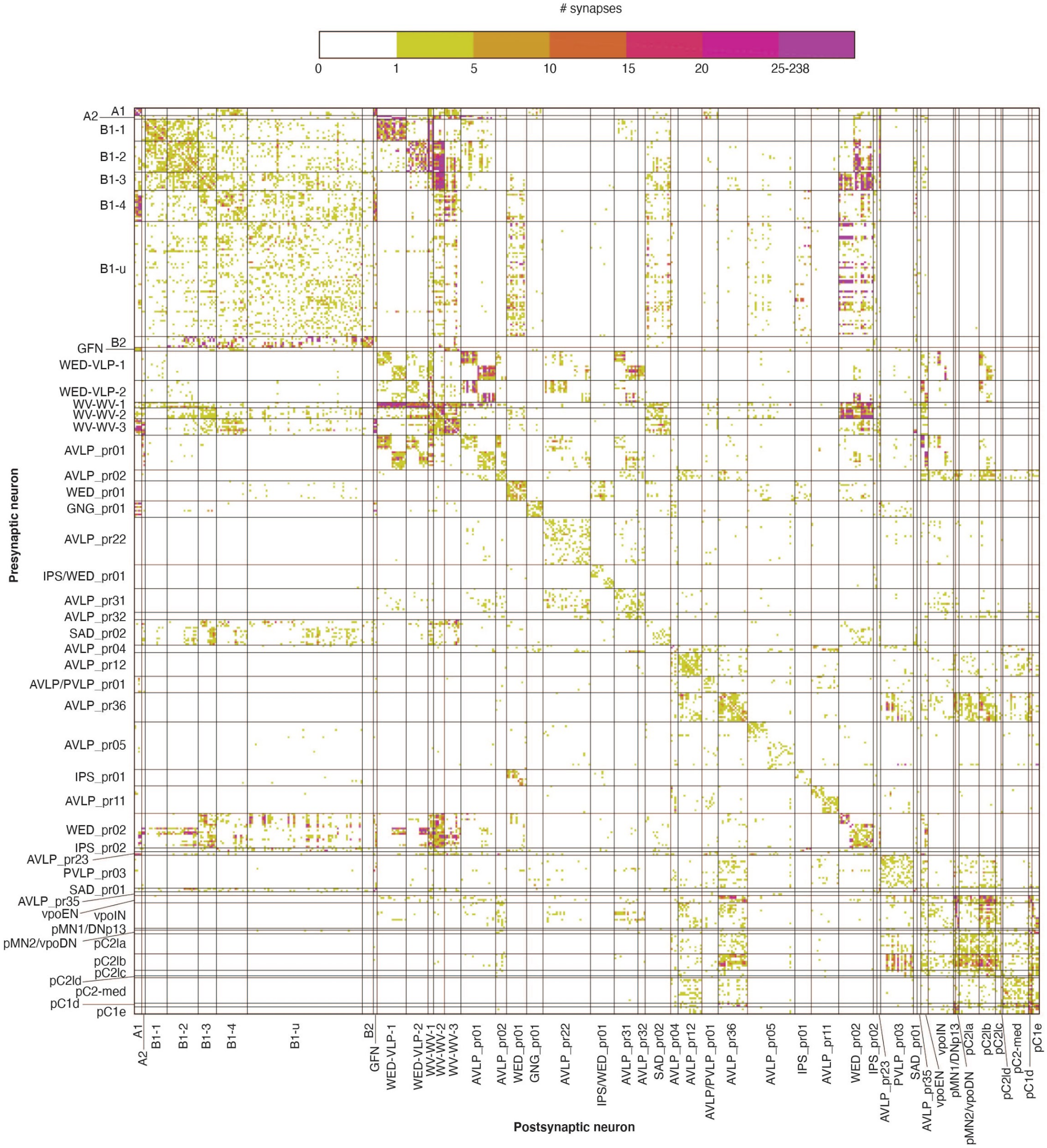

**Supplemental Figure 8. Synaptic connections between each auditory cell type in FlyWire (both hemispheres).** The number of synapses between every pair of neurons included in our EM connectivity analysis (see Figure 7; n=496 neurons), and organized by cell type. Synapse counts are based on an automated dataset (Buhmann et al., 2019). We include neurons from both the left and right hemisphere of the brain, and neurons with somas in the left hemisphere are listed first in each cell type grouping. Previously characterized auditory neurons are separated into their cell types according to (Dorkenwald et al., 2020). Synapses are plotted with presynaptic partners on the y-axis and postsynaptic partners on the x-axis. The color scale indicates the number of synapses between individual neuron pairs.

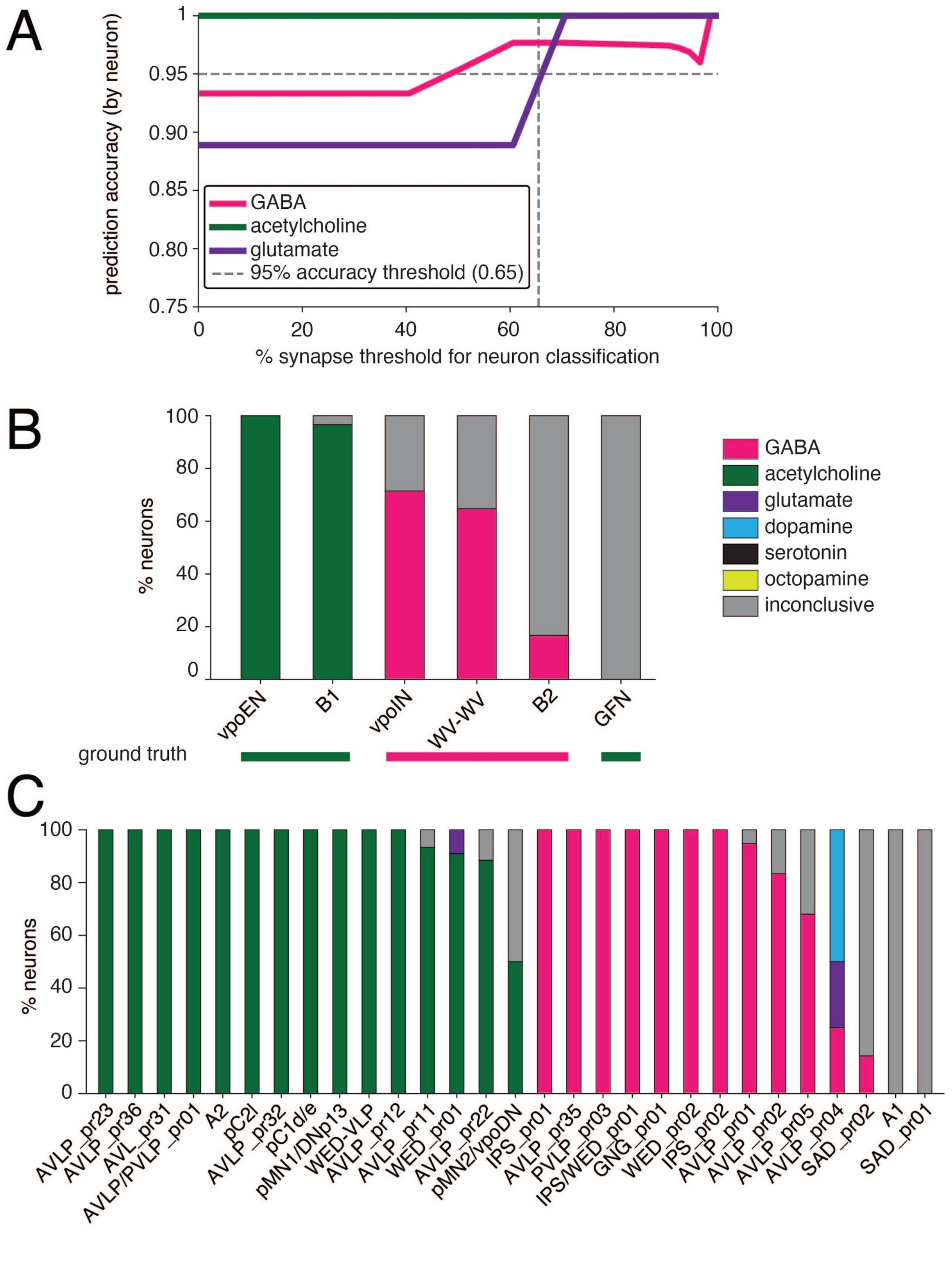
A
prediction accuracy (by neuron)
1
0.95
0.90
0.85
0.80
0.75
0
20
40
60
80
100
% synapse threshold for neuron classification
GABA
acetylcholine
glutamate
95% accuracy threshold (0.65)
B
% neurons
100
80
60
40
20
0
vpoEN
B1
vpoIN
WV-WV
B2
GFN
ground truth
GABA
acetylcholine
glutamate
dopamine
serotonin
octopamine
inconclusive
C
% neurons
100
80
60
40
20
0
AVLP_pr23
AVLP_pr36
AVL_pr31
AVLP/PVLP_pr01
A2
pC2l
AVLP_pr32
pC1d/e
pMN1/DNp13
WED-VLP
AVLP_pr12
AVLP_pr11
WED_pr01
AVLP_pr22
pMN2/vpoDN
IPS_pr01
AVLP_pr35
PVLP_pr03
IPS/WED_pr01
GNG_pr01
WED_pr02
IPS_pr02
AVLP_pr01
AVLP_pr02
AVLP_pr05
AVLP_pr04
SAD_pr02
A1
SAD_pr01

**Supplemental Figure 9. Neurotransmitter predictions for auditory cell types from EM.** A) Prediction accuracy as a function of the fraction of synapses that have been predicted as the respective majority neurotransmitter. Data come from a test set of 135 neurons in a prior study (Eckstein et al., 2020). A threshold of 65% of consensus synaptic neurotransmitter predictions across a neuron's synapses results in greater than 95% classification accuracy for the three "classical" neurotransmitters. We used this threshold for predicting the neurotransmitters of auditory neurons. B) We compared neurotransmitter predictions for six cell types with their classifications from the literature based on fluorescence in-situ hybridization or antibody staining (Allen and Murphey, 2007; Tootoonian et al., 2012; Lai et al., 2012; Wang et al., 2020b). The deep learning-based classifier predictions for each neuron of a cell type (in the FAFB EM volume) are shown in the histogram, and the ground truth results from the literature are shown below. Neurons for which less than 65% of synapses are predicted to have the same neurotransmitter are labeled as "inconclusive" (grey). C) Classifier predictions for each cell type shown in Figure 7. The neurotransmitters used by these cell types have not been reported previously.

Supplemental Figure 10

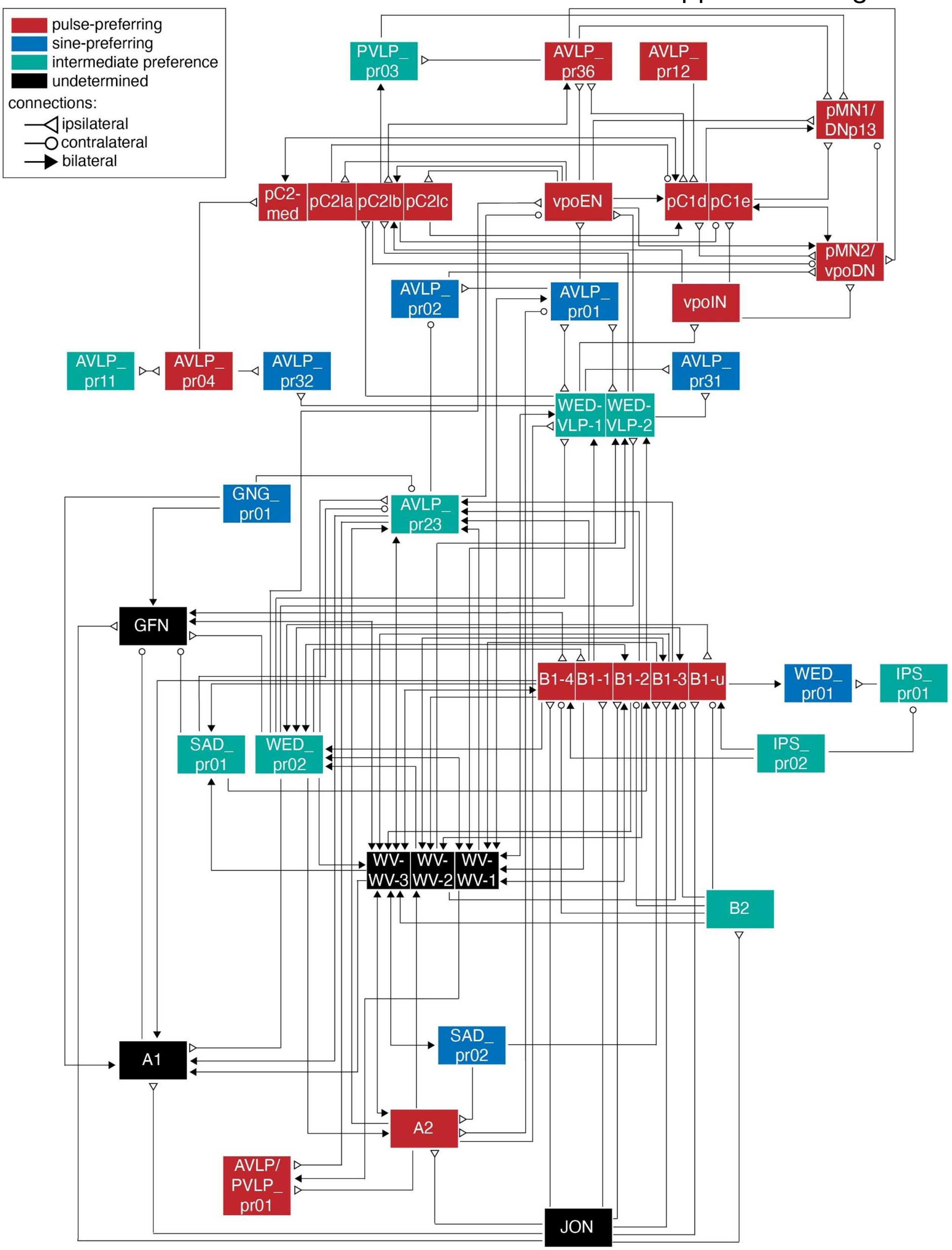

**Supplemental Figure 10. Auditory connectome with previously identified auditory neurons divided into subtypes.** Cell types B1, WV-WV, WED-VLP, pC2, and pC1 are broken down by their known subtypes (Dorkenwald et al., 2020; Wang et al., 2020a; Deutsch et al., 2020; Xu et al., 2020). The song mode preferences of these cell types comes from recordings of GAL4 lines that typically label multiple subtypes (or the broader cell class).

Supplemental Figure 11

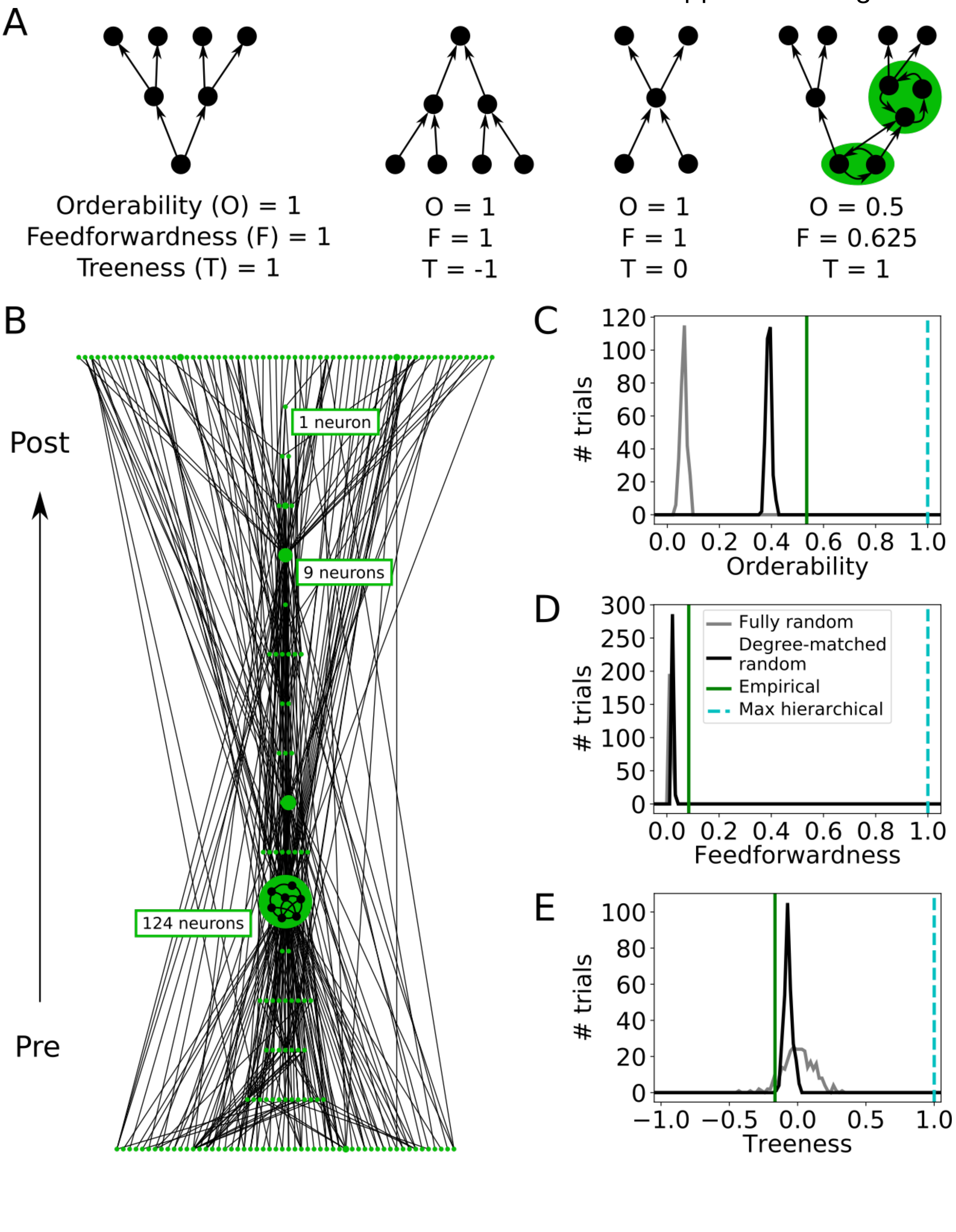

**Supplementary Figure 11: Evaluating the hierarchical structure of the auditory network**. A. Measures of hierarchical structure (orderability, feedforwardness, and treeness) for four simple networks (reproduced from [Corominas-Murtra et al., 2013]); each green region highlights a strongly connected component (a set of neurons all mutually reachable from one another via at least one directed path). B. Hierarchically displayed node-weighted graph condensation of the actual auditory network. Each node in the condensation is a strongly connected component of the original auditory network, with node size indicating the number of neurons in the component (minimum 1 neuron, maximum 124 neurons). Connections are oriented upward with postsynaptic targets displayed above presynaptic sources. C. Orderability of the auditory network, computed from the original network and the graph condensation (A), as compared to 300 instantiations (trials) of either a fully random network (with only neuron count and connection probability matched to the original auditory network) or degree-matched random network (in which each neuron's incoming and outgoing connection numbers were inherited from the empirical network but connections were otherwise randomized). D. Feedforwardness of the auditory network, as compared to fully random and degree-matched random networks. The low value arises from most paths passing through the largest connected component. E. Treeness of the auditory network, as compared to fully random and degree-matched random networks. Prior to analysis, connections between two neurons in the original network were only counted if at least 15 synapses were detected (171 out of 496 neurons were unconnected to the main network and left out of the calculations). A maximally hierarchical network has a feedforwardness, orderability, and treeness of 1 (cyan; C-E).
